# Fluid-structure interaction analysis of eccentricity and leaflet rigidity on thrombosis biomarkers in bioprosthetic aortic valve replacements

**DOI:** 10.1101/2022.01.06.475272

**Authors:** David Oks, Mariano Vázquez, Guillaume Houzeaux, Constantine Butakoff, Cristóbal Samaniego

**Affiliations:** Barcelona Supercomputing Center (BSC), Barcelona, Spain; ELEM Biotech, Barcelona, Spain

## Abstract

This work introduces the first 2-way fluid-structure interaction (FSI) computational model to study the effect of aortic annulus eccentricity on the performance and thrombogenic risk of cardiac bioprostheses. The model predicts that increasing eccentricities yield lower geometric orifice areas (GOAs) and higher normalized transvalvular pressure gradients (TPGs) for healthy cardiac outputs during systole, agreeing with *in vitro* experiments. Regions with peak values of residence time and shear rate are observed to grow with eccentricity in the sinus of Valsalva, indicating an elevated risk of thrombus formation for eccentric configurations. In addition, the computational model is used to analyze the effect of varying leaflet rigidity on both performance, thrombogenic and calcification risks with applications to tissue-engineered prostheses, observing an increase in systolic and diastolic TPGs, and decrease in systolic GOA, which translates to decreased valve performance for more rigid leaflets. An increased thrombogenic risk is detected for the most rigid valves. Peak solid stresses are also analyzed, and observed to increase with rigidity, elevating risk of valve calcification and structural failure. The immersed FSI method was implemented in a high-performance computing multi-physics simulation software, and validated against a well known FSI benchmark. The aortic valve bioprosthesis model is qualitatively contrasted against experimental data, showing good agreement in closed and open states. To the authors’ knowledge this is the first computational FSI model to study the effect of eccentricity or leaflet rigidity on thrombogenic biomarkers, providing a novel tool to aid device manufacturers and clinical practitioners.

## 1 Introduction

Heart valve disease represents a critical clinical problem worldwide. Its prevalence is increasing globally due to degenerative valve disease in aging populations in developed countries, and due to inadequate treatment of rheumatic heart disease in the developing world. As a result 130,647 aortic valve replacement (AVR) procedures were carried out in 2019 in the United States alone [1]. Despite the long-term durability of mechanical aortic valve replacements (MAVRs), they require life-long administration of anticoagulants and are implanted invasively. In contrast, bioprosthetic aortic valve replacements (BAVRs) don’t require anticoagulants, but they have a limited durability, and typically fail 10–15 years post implantation, making MAVRs the prosthesis of choice for patients below 60 years of age. BAVRs are constructed from porcine heart valves or bovine pericardium preserved with glutaraldehyde, and may be surgical (SAVR) or transcatheter (TAVR), with the latter delivered using a non-invasive procedure. The non-invasiveness and reduced thrombogenic risk of TAVRs explains its rapid adoption and approval for low-risk patients, accompanied by a reduction in the recommended minimum age for implantation [2, 3]. This has led the number of implanted BAVRs to surpass that of MAVRs in 2019. This work is thus focused specifically on the performance analysis of BAVRs.

Despite its fast adoption, the widespread use of TAVRs is limited by the evidence of subclinical leaflet thrombosis post-TAVR [4, 5, 6], which has been suggested as the underlying reason for hypoattenuated leaflet thickening, leading to reduced leaflet motion [7, 8]. Thrombosis is the pathological clotting of blood within a vessel. Two different mechanisms may lead to thrombus formation:

1. coagulation cascade of protein activation at low shear-rates (*<* 50s^−1^) generating erythrocyte-rich thrombi that cause them to have a red appearance [9, 10], and
2. cellular platelet aggregation at high shear rates (*>* 5000s^−1^) acting separately or in concert to prevent excessive bleeding in a process called hemostasis, producing white clots [11, 12].

In either type, a piece of thrombus may break off as an embolus, traveling through the circulation and lodging somewhere else causing a vascular occlusion, ischemia, and eventually death. Often artificial surfaces such as those of prosthetic heart valves may initiate thrombosis through the contact activation pathway of coagulation. Low flow rate and regions of flow stagnation near the valve have been suggested as the source of leaflet thrombosis since they have been observed in *in vitro* and *in silico* studies [13, 14, 15, 16, 17, 18]. Identifying these regions is crucial to assess the thrombogenic potential of BAVRs and improve designs, but determining them in experimental studies or clinical measurements has proven extremely challenging.

Computational modeling and simulation provides a powerful cost- and time-efficient tool to access detailed information of the flow and structure which is not visible in neither clinical imaging nor *in vitro* bench testing. Though extensive bibliography exists for *in vitro* studies, *in silico* analyses of thrombogenic potential are still mainly restricted to ventricular assist devices [19, 20] and aneurysms [21]. Mayo et al. analyzed the difference between two TAVR models on five different quantities of interest (QoIs) related to thrombogenic potential [22]. Their study consists of two phases: (a) A purely computational solid mechanics (CSM) model to deploy the stents and leaflets, followed by (b) a purely computational fluid dynamics (CFD) model to obtain the thrombosis QoIs for each TAVR, modeling the prostheses as rigid and fixed. It provides useful insight on how to quantify thrombogenic risk of TAVRs. On the other hand, authors do not account for the valve deformation or the 2-way fluid-structure interaction (FSI) during the pulsatile cycle. In the present work a fully 2-way FSI computational model is used to capture the pulsatile dynamics of fluid and solid domains. Biomarkers (*ie:* residence time and strain rate) are analyzed to assess the thrombogenic risk of bioprostheses. In order to understand calcification and dysfunction risk, an analysis is carried out on precise measures of stresses exerted on native and bioprosthetic tissue along the cardiac cycle. This model therefore provides a valuable complement to bench testing by enabling an in-depth assessment of device performance, in a broad range of working conditions.

Another issue which worries both patients and clinicians is that of native and bioprosthetic valve calcification. Mechanical stress has been extensively related to calcification and structural failure of aortic valve prostheses [23, 24, 25]. Deiwick et al. show that leaflet tensile stress and shear stress may lead to valvular structural failure [24]. Moreover, Thubrikar et al. report that high compressive stress is correlated with the leaflet calcification of aortic valve prostheses [25]. Leaflet stresses are particularly difficult to measure in an experimental and/or clinical setup. An indirect form of measurement can be carried out *in vitro* by tracking strains [26]. Nonetheless this process is complex and prone to introduce multiple sources of error. Computational solid mechanics provides a means of accessing detailed information on solid strains and stresses on both surfaces and in volume. In addition, cardiac valve dynamics involve a highly nonlinear coupling between the fluid and the solid structure, therefore making it fundamental to model the full FSI to capture realistic leaflet stress distributions in predictive models. In this work, leaflet von Mises stresses are computed to analyze the calcification risk of bioprostheses.

Although some degree of eccentricity of the aortic root is observed in all patients [27], BAVRs are designed circular. For patients with aortic stenosis, heavy calcium deposition on the leaflets and the aortic root cause distortion of the post-intervention TAVR geometries, resulting in elliptical shapes [28, 29, 30] and causing paravalvular leakage [31]. Large degrees of eccentricity may affect leaflet coaptation and produce intravalvular regurgitation [29] and larger systolic pressure gradients [32] thus worsening prosthesis function. Moreover, without proper leaflet apposition, uneven distribution of stress on the leaflets may also affect long-term valve durability [33]. Although extreme aortic root eccentricities have been shown to produce adverse effects on prostheses performance, available studies present conflicting results: an *in vitro* analysis on the effect of different degrees of oversizing and eccentricity on TAVR performance found that effective orifice areas (EOAs) were larger and TPGs were lower for elliptical compared to circular annuli [34], concluding that the performance was improved with slight degrees of eccentricity. Realistic computational models provide a useful tool to shed some light on this yet unclear subject.

Some computational studies have been carried out to study this parameter in the past. In Sun et al. an idealized TAVR geometry in a calcified aortic root was used to study the effect of varying eccentricity on stress distribution and valve leakage in open and closed states [35]. The valve structure was modeled as rigid, using a purely CFD approach, therefore not capturing the non-linear FSI feedback. Finotello et al. analyze the effect of different purely structural finite element modeling strategies of a TAVR deployment on QoIs, such as eccentricity of the deployed TAVR [36]. Results are compared to post-operative computational tomography scans of real implants. Sirois et al. study the effect of eccentric deployment and under-expansion of TAVRs hydrodynamic performance (such as on turbulent viscous shear stress) by using rigid models for the prostheses structure in an open state [37]. They observed that only extreme eccentricities affect TAVR performance, as does an under-expansion of the stent. A realistic fully-FSI computational model is long overdue to assess the effect of a wide range of working conditions on physiologically relevant QoIs for this problem. In this work, a FSI model is used to analyze the effect of aortic root eccentricity on valve performance, thrombogenic and calcification risk in pathological, healthy rest and healthy exercise working conditions.

In addition to aiding the design process, verified and validated software may be employed as additional evidence to support claims of device safety and effectiveness in regulatory submissions [38]. A further step is to use *in silico* models to plan clinical interventions and aid diagnoses, minimizing patient risk. For clinical applications, models should be capable of adapting to patient-specific conditions. In order for *in silico* models to be employed in both regulatory and clinical applications, Verification and Validation (V&V) is critical to prove credibility of the software, as explained in ASME’s V&V40 guidelines [39]. In the following the FSI method is validated against a well known numerical FSI benchmark [40]. The ISO 5840-3 standard imposes certain requirements on the transvalvular pressure gradient (TPG) and EOA of valve prostheses [41]. In this work these QoIs are computed and qualitatively contrasted against analogous *in vitro* experiments [42, 32] as a first step in the V&V of the introduced computational model.

To the authors’ knowledge there is no published computational model which analyzes the effect of either rigidity or eccentricity of the aortic root on TAVR performance and thrombogenic risk while accounting for the full FSI between blood flow and the bioprosthesis. In the following work, a parallel immersed two-way FSI computational scheme for unstructured meshes is implemented in *Alya*, Barcelona Supercomputing Center’s (BSC’s) in-house high-performance computing (HPC) multi-physics software. The method is introduced in section 2 and validated against the well-known FSI3 benchmark of Turek & Hron [40] in section 3.1. In sections 3.2 and 3.3 the effect on BAVR performance is analyzed respectively for the following parameters:

1. leaflet rigidity, and
2. aortic annulus eccentricity.

For each of these parameters, in sections 3.2 and 3.3 respectively, the following QoIs are used to evaluate the valve performance according to the ISO 5840-3 standard [41], and to contrast the model against the experimental data:

1. transvalvular pressure gradients, and
2. geometric orifice area.

In each section, the effect of either leaflet rigidity (section 3.2) or eccentricity (section 3.3) on thrombogenic risk is quantified by evaluating the following thrombosis biomarkers:

3. residence time, and
4. shear rate.

In each case, the risk of calcification and structural failure is also evaluated with the

6. von Mises stresses.

In section 4 conclusions are given. Finally, limitations of this work and a roadmap for the future of *in silico* modeling of cardiac valve bioprostheses are presented in sections 4.1 and 4.2 respectively.

## 2 Methods

In this section, the governing equations and numerical discretizations are presented for the fluid and solid mechanics problems. Next, relevant issues on FSI numerical methods are briefly discussed, and the immersed finite element method used in this work is presented. Finally, relevant aspects of the parallel computational implementation are explained.

### 2.1 Fluid mechanics

In this work, blood is modeled as a Newtonian incompressible fluid. The Newtonian approximation is an acceptable assumption and not far from reality in a significant part of the circulatory system under normal conditions. This is specially true in large blood vessels where suspended particles (*ie:* red blood cells) are well below the characteristic sizes of the vessels (*ie:* left ventricle outflow tract, aortic root and aorta) [43]. Hence haemodynamics are modeled in this work using the incompressible Navier-Stokes equations, which are formulated below in the Eulerian frame of reference.

#### 2.1.1 Continuum form of Navier-Stokes equations

For a fluid moving in the domain Ω^*f*^ bounded by Γ^*f*^ = *∂*Ω^*f*^ during the time interval (*t*_0_, *t*_*f*_), the Navier-Stokes equations consist in finding a velocity ***u*** and kinematic pressure *p* such that:

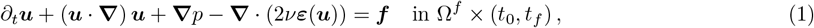

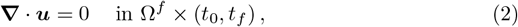

where *ν* is the kinematic viscosity, 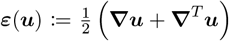 is the velocity strain rate tensor and ***f*** is the vector of external body forces. The Navier-Stokes equations are supplemented by Dirichlet boundary conditions which prescribe the velocity, and Neumann boundary conditions which prescribe the traction 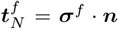, with ***σ***^*f*^ =−*p****I*** + 2*μ****ε***(***u***) the Cauchy stress tensor and ***n*** the outward normal vector to the surface Γ^*f*^. Let 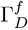 and 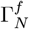 be the Dirichlet and Neumann parts of the boundary Γ^*f*^ respectively, such that 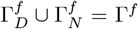 and 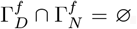. The boundary conditions consist in prescribing

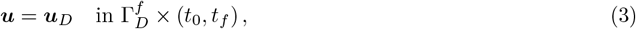

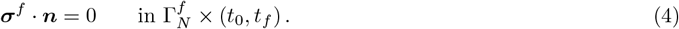

Additionally, initial conditions must be set on the problem:

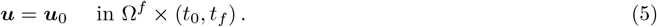

To obtain the weak form of the Navier-Stokes equations (1–2), the spaces of vector functions 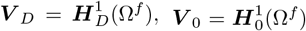 and *Q* = *L*^2^(Ω^*f*^) are introduced. Here *L*^2^(Ω^*f*^) is the space of square-integrable functions, *H*^1^(Ω^*f*^) is a subspace of *L*^2^(Ω^*f*^) formed by functions whose derivatives also belong to 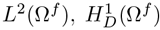 is a subspace of *H*^1^(Ω^*f*^) that satisfies the Dirichlet boundary conditions on 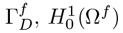 is a subspace of *H*^1^(Ω^*f*^) whose functions are zero on Γ^*f*^, and 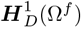 and 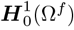 are their vector counterparts in the 3-dimensional Euclidean space ℝ^3^. In the following (*·, ·*) indicates the standard *L*^2^ inner product.

For the dynamic case *V*_*t*_ ≡ *L*^2^(*t*_0_, *t*_*f*_; *V*_*D*_) and *Q*_*t*_ ≡ *D*′(*t*_0_, *t*_*f*_; *Q*) are introduced, where *L*^*p*^(*t*_0_, *t*_*f*_; *X*) is the space of time dependent functions in a normed space Σ such that 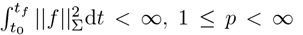 and *Q*_*t*_ consists of mappings whose *Q*-norm is a distribution in time. The weak form of problem (1-2) with the boundary conditions defined in (3-4) is then: find ***u*** ∈ ***V*** _*t*_ and *p* ∈ *Q*_*t*_ such that:

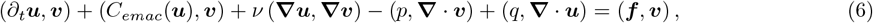

for all (***v***, *q*) ∈ ***V*** _0_*×Q*. In this method the nonlinear term is written using the energy, momentum and angular momentum conserving form (EMAC):

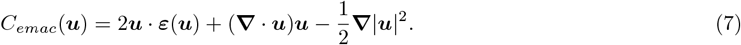

#### 2.1.2 Space discretization of Navier-Stokes equations

The spatial discretization is constructed using the finite element (FE) method. The discrete linear subspaces that approximate the respective continuous spaces defined above are ***V*** _*Dh*_ ⊂ ***V*** _*D*_, ***V*** _0*h*_ ⊂ ***V*** _0_ and *Q*_*h*_ ⊂ *Q*. Equal (linear) interpolation is used for both velocity and pressure. The space-discretized problem reads: find 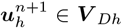 and 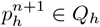 such that:

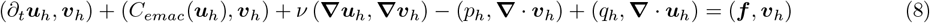

for all virtual (***v***_*h*_, *q*_*h*_) ∈ ***V*** _0*h*_ *× Q*_*h*_. These equations can then be written in the following matrix form:

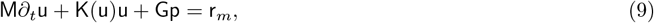

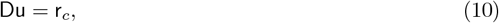

where `u` and `p` are the arrays of the nodal unknowns for ***u*** and *p* respectively. Denoting the nodal indices with superscripts *a, b*, the space indices with subscripts *i, j* and the linear shape functions of node *a* as *N*^*a*^, the matrices involved above are:

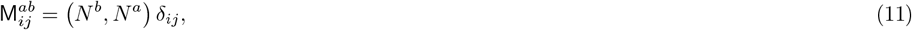

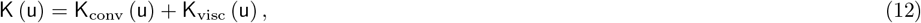

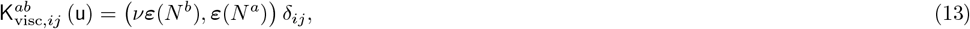

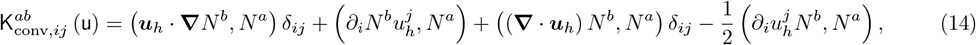

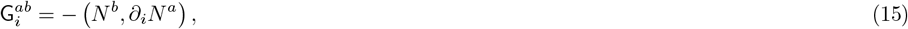

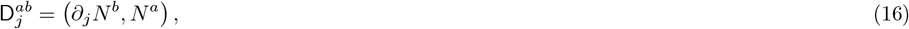

where *δ*_*ij*_ is the Kronecker delta, and 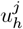 notes the *j*^th^ component of vector `u`_*h*_. The vectors `r`_*c*_ and `r`_*m*_ include terms coming from the application of Dirichlet boundary conditions and the latter also includes the contribution from the body forces. Since here an explicit time discretization is used, the mass matrix is lumped to avoid the solution of a linear system for the velocity as is usually done for explicit schemes in FE methods. In the following, M refers to the lumped mass matrix. Defining the vector `B` = −`Ku` + `r`_*m*_, equation (9) can be re-written as

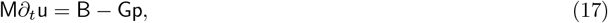

which facilitates the derivation of the time discretization.

#### 2.1.3 Time discretization of Navier-Stokes equations

An explicit Runge-Kutta is used to time-discretize equations (17) and (10), imposing a divergence-free constraint with a fractional step scheme. This implies solving the following equations:

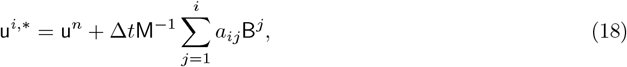

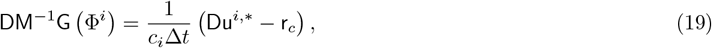

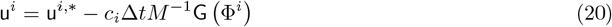

for substeps *i* = 2, …, *s* (for the first substep *u*^1^ = *u*^*n*^) and finally obtaining the unknowns at the new step from

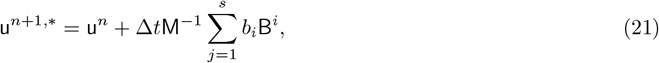

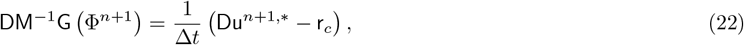

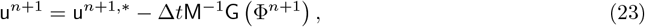

where *a*_*ij*_, *b*_*i*_ and *c*_*i*_ = Σ_*j*_ *a*_*ij*_ are the coefficients and *s* the number of substeps of the Runge-Kutta scheme respectively, Δ*t* is the time step size, and the pseudo-pressure Φ is a first order approximation to the pressure (*ie:* Φ = *p* + 𝒪 (Δ*t*)). To reduce computational cost, the discrete Laplacian *DM*^−1^*G* is approximated as:

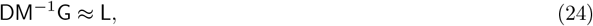

which introduces a stabilizing effect for the pressure that allows to use equal order interpolation, while only introducing an error of the same order as the pressure interpolation [44]. For more details on this low dissipation FE incompressible Navier-Stokes formulation, refer to Lehmkuhl et al. [45].

### 2.2 Solid mechanics

In computational solid mechanics (CSM), meshes are usually described as Lagrangian, that is, in the reference system fixed to the moving and/or deforming solid. Within Lagrangian FE descriptions two approaches are commonly taken:

1. **Total Lagrangian formulation:** derivatives and integrals are taken with respect to Lagrangian (material) coordinates ***X***^*s*^.
2. **Updated Lagrangian formulation:** derivatives and integrals are taken with respect to Eulerian (spatial) coordinates ***x***^*s*^.

In this work total Lagrangian formulation is used. To describe it properly, it is necessary to define the transforms between reference systems.

#### 2.2.1 Continuum solid mechanics equations

Given an arbitrary deformable body 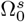 in a 3-dimensional Euclidean space ℝ^3^, the position of a material point of the body at the reference time *t* = *t*_0_ is described by the vector ***X***^*s*^, while its position at the current time *t* ∈ [*t*_0_, *T*] is given by the vector ***x***^*s*^. The current position vector is the image of the regular map ***χ***(***X***^*s*^, *t*) which describes the motion of the body. The displacement field is defined as

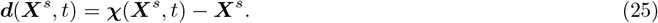

Assuming the body is subjected to body forces ***b***, the governing equations can be written in the total Lagrangian form as:

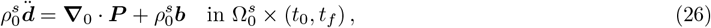

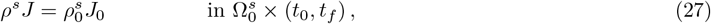

corresponding to the conservation of linear momentum and mass respectively, in the reference (undeformed) configuration. Here *ρ*^*s*^ and 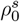 are the solid’s mass density in the current and reference configurations respectively, 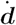 and 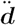 represent the velocity and acceleration respectively, 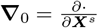. indicates the gradient operator with respect to the reference coordinates, ***P*** is the nominal stress tensor, and *J* = det(***F***), with the deformation tensor defined as 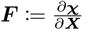. The governing equations must be supplemented with boundary conditions. Let 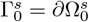 denote the outer boundary of the domain, with outward vector ***n***. Dirichlet boundary conditions are imposed on 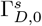 and Neumann boundary conditions on 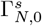, such that 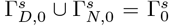 and 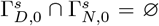. These correspond to imposing displacements ***d***_*D*_ and tractions 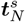 respectively:

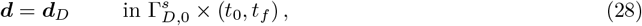

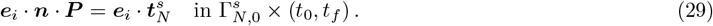

The system must be appended with initial conditions:

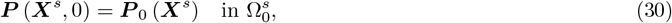

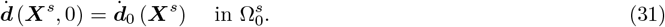

To complete the description, a constitutive equation must be given for the material. Elastic materials for which the work is independent of the load path are said to be hyperelastic. These are characterized by the existence of a strain energy function *ψ* that is a potential for the second Piola-Kirchoff stress ***S***:

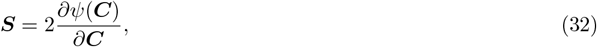

where ***C*** = ***F***^*T*^ · ***F*** is the Cauchy Green strain tensor. The nominal stress can then be derived from the second Piola-Kirchoff tensor as

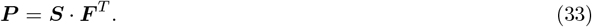

Note that the first Piola-Kirchoff stress tensor is the transpose of the nominal stress tensor, *PK*1 = ***P*** ^*T*^. Bovine and porcine pericardium which are the most frequent materials used to confection bioprosthetic heart valve replacements are formed by an isotropic extracellular matrix and an anisotropic alignment of collagen fibers. This anisotropy may be modeled with fiber-reinforced anisotropic hyperelastic materials [46, 47]. Nonetheless, material anisotropy is out of the scope of the current work. The validity of this simplification is supported by the fact that the present aortic valve model is compared to an experiment in which an isotropic polyurethane valve prosthesis was used. Therefore in the remainder of this work the Neo-Hookean material model is used for the solids involved. This model is an extension of Hooke’s law to large deformations. The free energy *ψ* for a Neo-Hookean material can be expressed as:

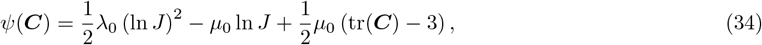

where *λ*_*s*_ and *μ*_*s*_ are known as the Lamé coefficients. The second Piola-Kirchoff stress tensor is then given by

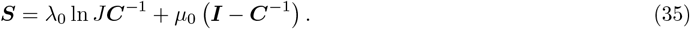

To derive the weak form of the momentum balance equation, the spaces of vector functions 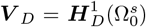 and 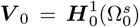 are introduced. An abuse of notation is done here by using the same notation as for the fluid domain. Once gain, for the dynamic case the subspace *V*_*t*_ ≡ *L*^2^(*t*_0_, *t*_*f*_; *V*_*D*_) is defined. As usual, the weak form of the momentum balance equation (26) consists of finding the displacements ***d*** ∈ ***V*** _*t*_ such that

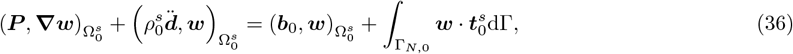

for any virtual displacement ***w*** ∈ ***V*** _0_. Here 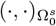 denotes the *L*^2^ norm integrated in the reference configuration 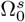.

#### 2.2.2 Spatial discretization of solid equations

As for the fluid, the reference continuum body 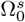 is discretized using a FE approximation. Let 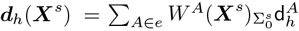 be the polynomial approximation of degree *k* (in this case *k* = 2) of the actual displacement ***d***, where *A* notes the nodal indices, d_*h*_ are the nodal values of displacement, and the FE shape functions for the solid are noted as *W* (***X***^*s*^). Hence, the matrix form of (26) reads

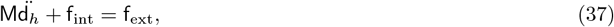

where `M, f`_int_ and `f`_ext_ are the mass matrix, internal and external force vectors respectively which write

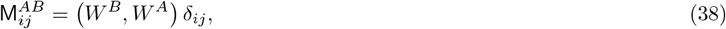

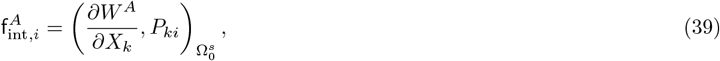

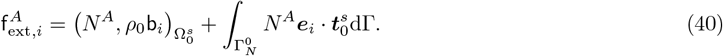

Note that all expressions are written in the reference configuration.

#### 2.2.3 Time discretization of solid equations

To discretize (37) in time, the generalized Newmark formulation is considered at time steps *t*^*n*^ and *t*^*n*+1^:

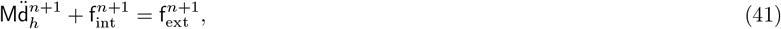

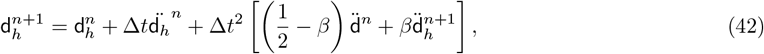

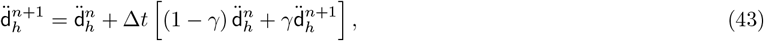

where parameters *β* and *γ* set the characteristics of the Newmark scheme. Parameter *β* = 0 results in an explicit scheme, whereas values of 0 *< β* ≤ 0.5 yields an implicit scheme, which is the option chosen in this work. In the implicit scheme the set of equations (41–43) are solved for the unknown displacement 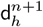, velocity 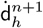 and acceleration 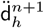 using the iterative Newton-Raphson algorithm.

### 2.3 Fluid-structure interaction

Heart valves consist of a soft-tissue of density very similar to that of blood. Their mechanics are therefore dictated by an intricate non-linear feedback interaction between structure and flow. This makes it mandatory to consider 2-way FSI models in order to correctly reproduce the dynamics for the full cardiac cycle. Moreover, this problem presents particular challenges in the realm of numerical methods: a near unity fluid-to-solid density ratio which produces instabilities in the fluid-structure coupling (*ie:* the *added-mass* instability), moderate Reynolds number flows (*ie: Re*; ≲ 6000) further favoring the instability of the system and large deformations of the fluid-solid interface. In numerical FSI methods the fluid mesh may adapt to the solid movement/deformation or it may be independent of the solid behavior as depicted in Figure 1. In this work, these families of methods are referred to as boundary conforming (BCMs) and non-boundary conforming methods (NBCMs) respectively. BCMs such as the arbitrary Lagrangian-Eulerian (ALE) method [48] yield accurate solutions near the fluid-structure interface. Nonetheless, with large displacements/deformations of the interface remeshing is required, incurring a very high computational cost. Moreover, for problems involving close contact, such as valve opening and closure, other issues such as grid element inversions may occur, further complicating the use of BCMs to simulate heart valves. In contrast, NBCMs introduce interpolation errors when transferring information between the overset fluid and solid meshes. However, since both meshes are independent, NBCMs are well posed when dealing with large displacements and deformations of the interface. The limitations in accuracy can be compensated by improved interpolation and coupling schemes. Consequently, NBCMs have been the preferred type of method for simulating heart valve dynamics since their inception [49]. With this in mind, this work introduces an immersed NBCM designed to couple unstructured FE discretizations of fluid and solid domains in problems involving large deformations of the fluid-solid interface.

**Figure 1:**
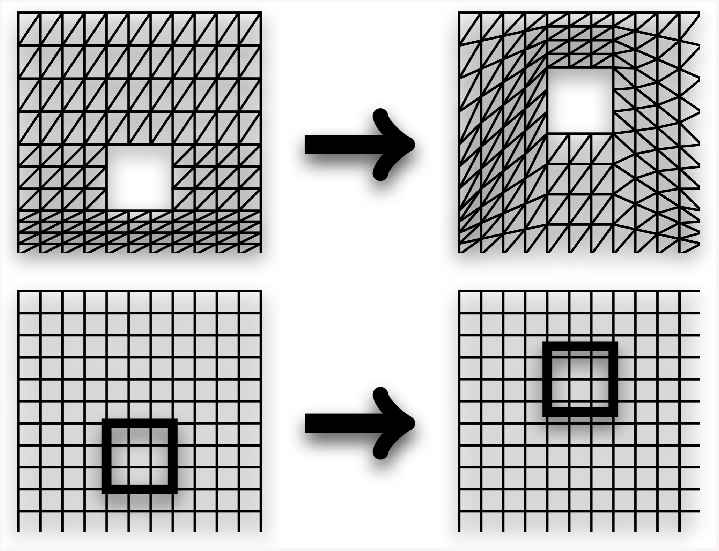
Sketch of BCM (top) and NBCM (bottom) under displacement of overlapping domains.

#### 2.3.1 Continuum form of FSI problem

In the immersed FSI coupling method presented here a body force term ***f***^*FSI,f*^ is added to the Navier-Stokes momentum equation (1) to account for the solid in this region. The force originates from the solid’s internal stresses produced due to the solid deformations. This term is computed in the Lagrangian solid domain and spread out from the Lagrangian to the Eulerian mesh. In the continuum domain this spreading operation can be described using Dirac delta distributions *δ*(***x***):

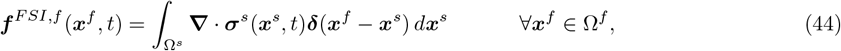

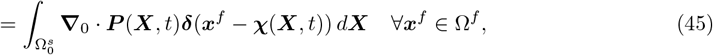

where the first line is given in the current configuration using the solid Cauchy stress tensor ***σ***^*s*^ = *J* ^−1^***F · P*** and the second in the reference configuration, using the nominal stress ***P***. On the other hand, the fluid velocities are interpolated to the Lagrangian solid mesh and solid nodes are directly displaced according to the interpolated velocities:

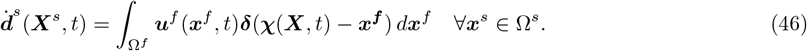

The Dirac delta distribution is defined such that for any compactly supported continuous function *f* (*x*),

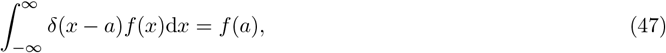

and the 3-dimensional delta function, ***δ***(***x***), is the product measure of the 1-dimensional delta functions in each variable separately:

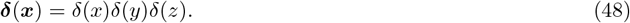

The Delta function is approximated by the interpolator *ϕ*(*x*) which should satisfy[50]: (1) continuity of velocities and forces across the fluid-solid interface, (2) the condition of completitude, and (3) reproducibility. In this work the FE interpolator is used to exchange information between the fluid and solid meshes. Wang and Zhang have shown that this interpolator satisfies all three conditions: continuity, completitude and reproducibility [50].

#### 2.3.2 Spatial discretization of FSI problem

On the discrete level, FE discretizations are used for both the fluid and solid domains. Therefore the FE interpolation operator is used to interpolate velocities from the fluid to the solid domain

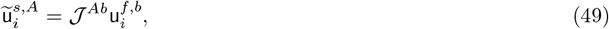

where the interpolation operator can be written in its discrete form as:

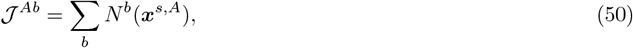

with *A* and *b* representing the solid and fluid nodal indices respectively and *N* (*x*) the fluid domain FE shape functions. Conversely, the transpose of the interpolation operator,

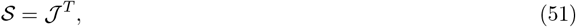

is used to spread forces from the solid to the fluid mesh:

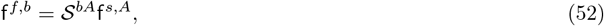

with 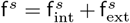 the sum of solid internal and external forces. It can be proven that 𝒮 conserves the total force. This interpolation scheme requires that the mesh size ratio between the background (fluid) and embedded (solid) meshes in the region of interaction must be close to 2:1, which is respected in the following work.

#### 2.3.3 Time discretization of FSI problem

Regarding the time-discretization, an explicit staggered time-stepping scheme is considered and summarized in Algorithm 1. Given the solution at time *t*^*n*^, the velocity `u`^*n*+1^ and pressure `p`^*n*+1^ at time step *t*^*n*+1^ = *t*^*n*^ + Δ*t* is obtained by solving the Navier-Stokes equations with the added volume force provided by the solid. The fluid velocities are then interpolated to the solid domain as *ũ*^*s,n*+1^, where the solid is deformed accordingly to these velocities using a 1st order forward Euler scheme to get *d*^*n*+1^ = *d*^*n*^ + *Δt* ũ^*s,n*+1^. Then solid internal forces 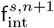 are computed in the deformed state. Finally the solid forces are spread out to the fluid mesh using operator 𝒮, and the procedure is repeated for each time step. To compensate the absence of FSI iterations in this staggered coupling scheme, a small time step size is considered (*ie:* Δ*t* ~ 20*μ*s). The time step is the same for both fluid and solid solvers, but the critical time step is dominated by the critical time step of the solid problem.

##### Algorithm 1 FSI coupling algorithm

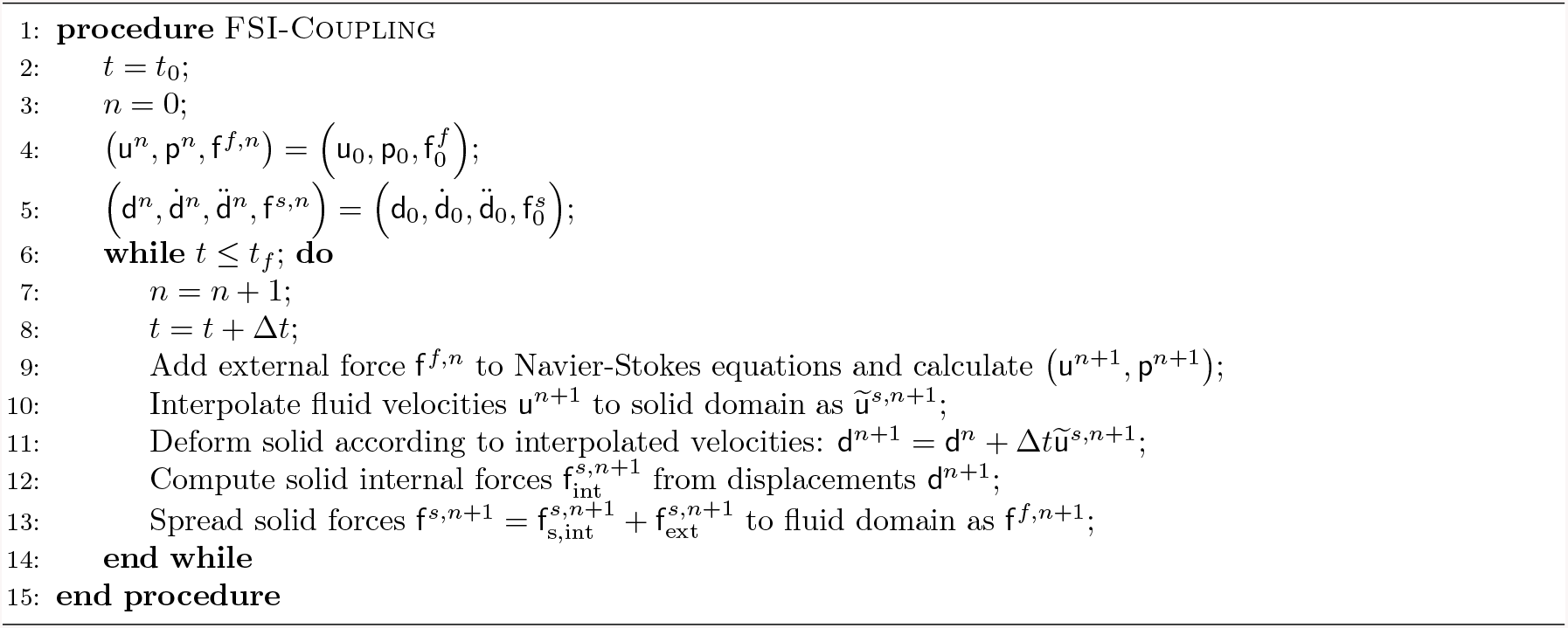

### 2.4 Computational aspects and parallel implementation

Heart valve mechanics involve an intricate and multi-physics interaction between complex deformable solids and blood flow in notably patient-specific conditions. This imposes challenging requirements on computational models, incurring in a very high computational cost. Running this type of simulations in practical times for both engineering and clinical applications requires the use of an efficient code capable of running in parallel on hundreds to thousands of processors.

#### 2.4.1 High performance computing

The present FSI model is implemented in *Alya*, BSC’s in-house multi-physics simulation software designed to run efficiently on supercomputers, exhaustively verified, validated, optimized and proved to give accurate solutions in complex fluid and solid mechanics problems, among other physics [51, 51, 52, 53, 54, 55, 45, 56]. *Alya* has been developed to scale efficiently in parallel on CPUs and/or GPUs using hybrid MPI, OpenMP, CUDA and/or

OpenACC models. It is one of the twelve simulation codes of the Unified European Applications Benchmark Suite (UEABS) and thus complies with the highest standards in HPC [57]. It therefore provides an ideal framework for the implementation of an AVR model required to run in practical times for engineering and clinical applications.

#### 2.4.2 Parallel implementation

A multi-code strategy is followed in this work, in which two instances of *Alya* are simultaneously executed (see Figure 2), CFD in one instance and solid mechanics (CSM) in another instance. At run time, each instance is partitioned in subdomains, being each subdomain pinned to an MPI task. In the cases analyzed here, the number of elements of the CFD problem is 200*×* larger than the CSM problem which is of a relatively small size given the thinness of the valve geometries (5.5 *×* 10^6^ elements for the fluid vs 25 *×* 10^3^ for the solid). Therefore, the nodal positions of all the solid domain are spread out to all CFD subdomains containing at least one solid node in their elements without significant loss in performance. The instances are then explicitly coupled in space as shown in Figure 2. The two problems are coupled in time using a staggered scheme in which fluid velocities are interpolated in volume to the solid nodes, and solid internal forces are spread out from the solid nodes to the fluid nodes of fluid elements containing at least one solid node.

**Figure 2:**
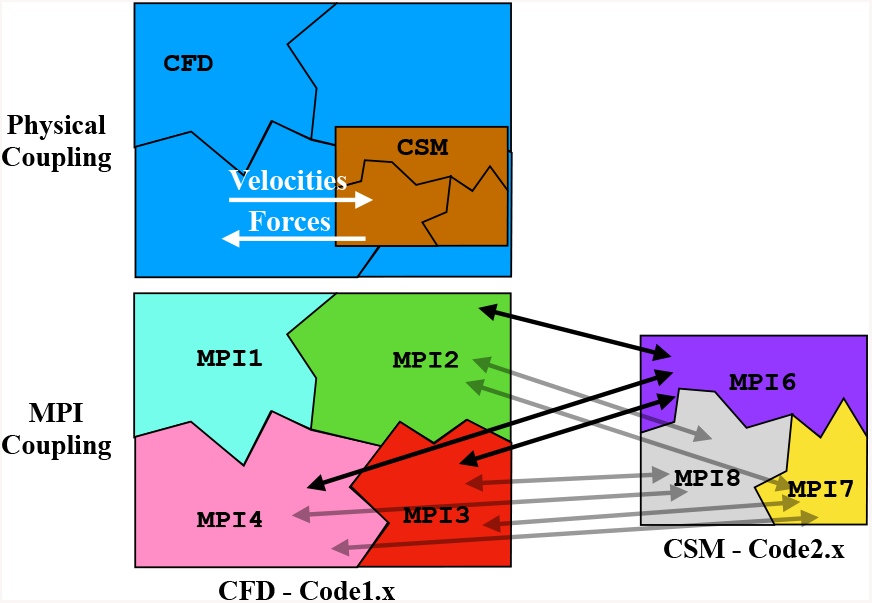
Parallel multi-code explicit coupling scheme between fluid and solid codes, the physical domain of each code is partitioned in multiple subdomains pinned to individual MPI tasks.

## 3 Results and discussions

The numerical FSI model is initially validated in section 3.1 against Turek & Hron’s 2-dimensional FSI3 benchmark used extensively in the literature [40]. In section 3.2 the numerical model is used to model a BAVR and qualitatively contrasted against the numerical-experimental work of Sigüenza et al. [42]. This same section analyzes the effect of the leaflet Young modulus on QoIs of valve performance as determined by the ISO 5840-3 standard, and thrombosis biomarkers employed in the literature. Finally, in section 3.3 the effect of annulus eccentricity on these standard QoIs and thrombosis biomarkers is studied and contrasted against experimental data from Kütting et al. [32].

### 3.1 Numerical validation: FSI3 benchmark

Turek & Hron’s FSI3 benchmark [40] has been used extensively to verify and validate multiple FSI computational models [56, 58, 59, 60, 61, 62, 63, 64]. This case consists of an elastic bar attached to a rigid circular hole, both embedded in a 2-dimensional Poiseuille flow as depicted in Figure 3. The nonlinear feedback between fluid and solid onsets the oscillations of the elastic tail as shown in Figure 4. Tip displacements (point A in Figure 3b) are recorded in time and their average values, amplitude and oscillation frequencies are then compared to bibliographic values. Fluid and solid densities are equal *ρ*^*f*^ = *ρ*^*s*^ = 1 g cm^−3^, while the fluid kinematic viscosity is *ν*^*f*^ = 10 cm^2^ s^−1^. The mean velocity is *Ū* = 2m s^−1^ from which the Reynolds number can be calculated as *Re* = *Ūd/ν*^*f*^ = 200. Finally the solid structure is modeled as an isolinear elastic material with Young modulus *E*^*s*^ = 5.6MPa and Poisson ratio *ν*^*s*^ = 0.4. As in Griffith et al. [64] this immersed FE method models solids as incompressible since velocities and therefore displacements are interpolated from the incompressible flow. Turek’s benchmark corresponds to a nearly-incompressible material, that is, *ν*^*f*^ = 0.4 (*ie:* the Poisson ratio *ν*^*f*^ ranges from 0 to 0.5 corresponding to the fully compressible and incompressible limits respectively). Therefore, differences between the current immersed FE method and reference values could be partially attributed to the incompressibility of the current solid model.

**Figure 3:**
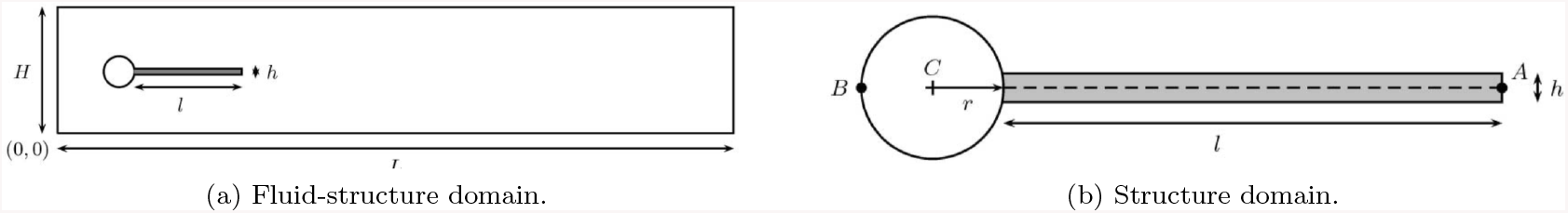
Domain for Turek & Hron’s FSI3 benchmark, diagram extracted from Turek & Hron [40].

**Figure 4:**
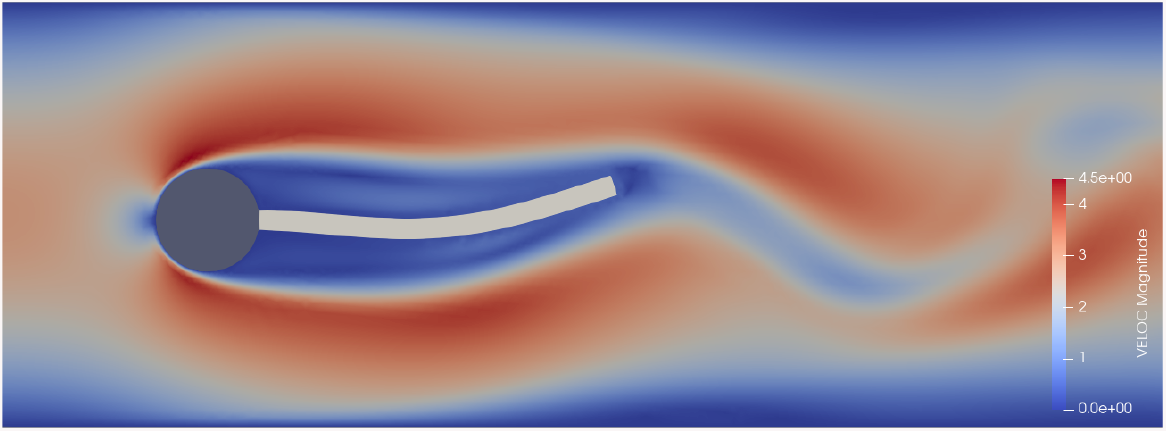
Snapshot of Turek FSI3 benchmark run used to validate the immersed FSI coupling in *Alya*.

A mesh convergence for this FSI3 benchmark is performed using both the ALE and immersed FSI methods implemented in *Alya*. Results for tip displacement time-average, amplitude and frequency are presented in Table 1 for each mesh resolution. Errors within 5% are observed for all quantities in the immersed method with respect to the reference values [40] for mesh resolutions equal to or below Δ*x* = 5mm, except for the mean displacement in *y*. Table 1 shows that errors are significantly reduced as mesh resolution is increased, showing the spatial convergence of the introduced algorithm. Moreover, it can be observed that errors are comparable and in some cases below those of the exhaustively validated ALE algorithm in *Alya*. Having verified the numerical consistency and validated the FSI model in a challenging transient 2-dimensional case, in the following section it is used to simulate a 3-dimensional deformable AVR.

**Table 1:** Results for the FSI3 benchmark [40] for both the immersed and ALE FSI coupling schemes implemented in *Alya*. The displacement of the center-right point of the beam (point A in Fig. 3b) is tracked in time. The columns correspond from left to right to: the FSI method used, the average element size, average tip displacement ⟨*d*_*i*_⟩, amplitude 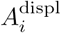 and frequency 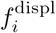 of oscillations for both horizontal and vertical directions, *i* = *x, y* respectively. The associated errors are computed with respect to reference values in Turek & Hron [40] and placed between parenthesis.

| Method | $\Delta x$ [mm] | $\langle d_x \rangle$ [mm] | $A_x^{\text{displ}}$ [mm] | $f_x^{\text{displ}}$ [Hz] | $\langle d_y \rangle$ [mm] | $A_y^{\text{displ}}$ [mm] | $f_y^{\text{displ}}$ [Hz] |
| --- | --- | --- | --- | --- | --- | --- | --- |
| Immersed | 10 | -0.19669 (-93%) | 0.074011 (-97%) | 6.9499 (-36%) | 2.3367 (58%) | 6.4033 (-81%) | 6.958 (31%) |
| Immersed | 5 | -2.7602 (2.6%) | 2.4017 (-5.1%) | 10.313 (-5.4%) | 1.9177 (30%) | 32.921 (-4.2%) | 5.1429 (-3%) |
| ALE | 10 | -2.3142 (-14%) | 2.0903 (-17%) | 10.872 (-0.26%) | 1.6055 (8.5%) | 29.397 (-14%) | 5.4367 (2.6%) |
| ALE | 5 | -2.356 (-12%) | 2.2025 (-13%) | 11.018 (1.1%) | 1.352 (-8.6%) | 30.146 (-12%) | 5.5086 (3.9%) |
| ALE | 2.5 | -2.4849 (-7.6%) | 2.2245 (-12%) | 10.872 (-0.25%) | 1.672 (13%) | 30.63 (-11%) | 5.4375 (2.6%) |
| ALE | 1.25 | -2.5469 (-5.3%) | 2.3508 (-7.1%) | 10.965 (0.6%) | 1.4332 (-3.2%) | 31.429 (-8.6%) | 5.4838 (3.5%) |

### 3.2 Effect of rigidity on aortic valve replacement

In this section an AVR FSI model is introduced, qualitatively contrasted against the numerical-experimental study by Sigüenza et al. in 2018 [42], and the effect of leaflet rigidity on valve performance is analyzed. In Sigüenza’s experiment a polyurethane heart valve model is placed in a rigid aortic chamber driven by a pulse duplicator. They prescribe a flat velocity profile at the inlet and a convective boundary condition at the outlet (*ie:* non-reflective outflow). As shown in Figure 5, the geometry of the experimental setup is approximated here, respecting the provided information: leaflet thickness, radius and boundary conditions. As described above, the leaflets are modeled using a Neo-Hookean material model. This section introduces an analysis of the effect of the Young modulus on standard QoIs of valve performance, and on biomarkers of thrombosis and leaflet calcification.

**Figure 5:**
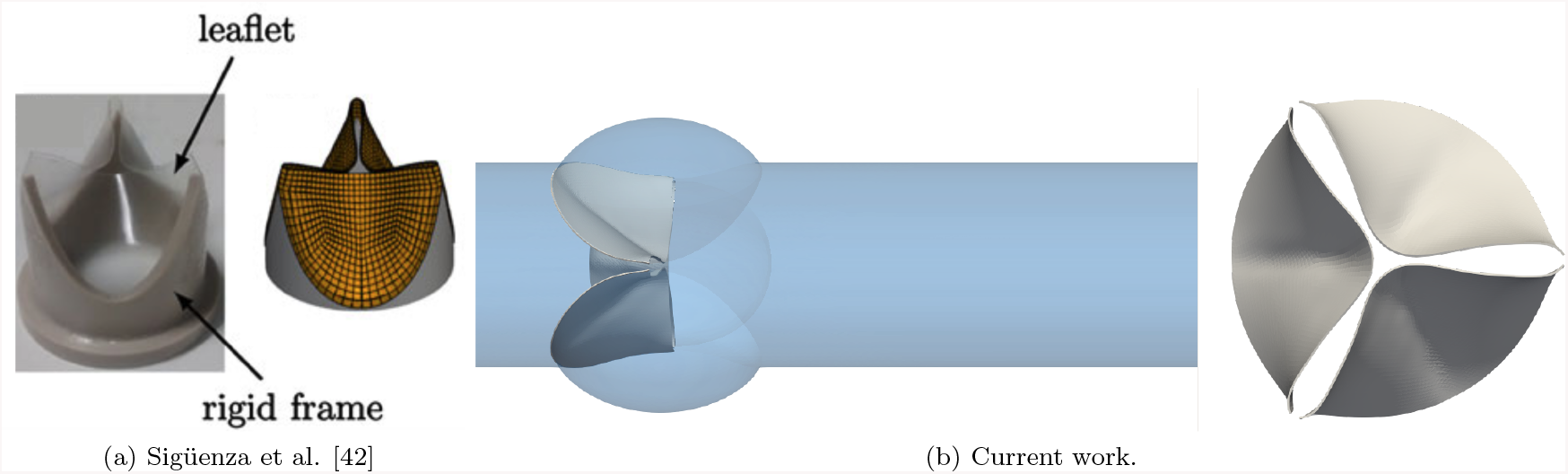
Geometries used for (a) experimental setup and numerical simulations in Sigüenza et al. [42] (figure adapted from this same publication), and (b) the current work numerical simulations contrasted against the reference [42].

The leaflet shear and bulk moduli given in the reference work[42], *G* = 2.4MPa and *K* = 1.6MPa respectively, yield a Poisson ratio of *ν* = 0.0. This value is far from realistic for polyurethane, a nearly incompressible material (*ie*: *ν*_polyurethane_ ~ 0.4 − 0.45). Given the uncertainty in the characterization of the polyurethane used, a parameter sweep is performed on the Young modulus, for values *E* ∈ *{*0.2, 0.5, 1.0, 2.0, 3.9*}*MPa, which serves three purposes:

1. To characterize the effect of leaflet rigidity on the performance tissue-engineered valve replacements.
2. To understand the effect of valve calcification on the performance of BAVRs by modeling calcification as an increase in leaflet rigidity.
3. To determine the sensitivity of the QoIs evaluated in this work to variations of the Young modulus, a parameter not clearly detailed in the reference work [42].

From this analysis we obtained the following main results:

1. The mean systolic orifice area decreases for more rigid leaflets, increasing the load on the left ventricle and potentially increasing the risk of left ventricular hypertrophy (LVH).
2. Systolic and (absolute) diastolic TPGs increase with leaflet rigidity, again elevating LVH risk.
3. Flow stagnation and peak shear rate regions become more localized for more rigid leaflets, leading to a higher thrombogenic risk.

#### 3.2.1 Experimental setup

The heart valve model was self-made by the authors of the reference work using a rigid frame made with PEEK material and leaflets manufactured out of thin polyurethane foil [42]. The valve was placed inside a rigid silicone aortic anatomy with a 25mm-diameter aortic root. The pulsatile flow is driven by a rotary pump which produces the periodic opening and closing of the valve as shown in Figure 6. The imposed flow waveform only considers the systolic phase and a subsequent backflow at the end of the ejection phase. A glycerol-water solution is used to model blood. Flow and structural parameters for the experiment are given in Table 2. Note that in the experimental setup a compliance is considered downstream of the valve, while this is not contemplated in the numerical model. Since neither compliance nor resistance parameters are not given, a compliance boundary condition is not included in this work (*ie:* Windkessel model [65]) at the outlet either.

**Table 2:** Parameters of experimental setup from Sigüenza et al. [42]

| Flow Parameters |  | Valve Parameters |  |
| --- | --- | --- | --- |
| Density | $\rho_f = 1.1 \text{ gcm}^3$ | Density | $\rho_f = 1.0 \text{ gcm}^3$ |
| Dynamic viscosity | $\mu_f = 3.6 \cdot 10^{-2} \text{ cP}$ | Shear modulus | $G = 2.4 \text{ MPa}$ |
| Heart rate | $n_{bpm} = 60 \text{ bpm}$ | Bulk modulus | $K = 1.6 \text{ MPa}$ |
| Mean cardiac output | $CO_{mean} = 3.48 \text{ Lmin}^{-1}$ | Leaflet Thickness | $e_l = 0.15 \text{ mm}$ |
| Reynolds number | $Re = 1388$ | Leaflet Radius | $R_l = 12.5 \text{ mm}$ |
| Womersley number | $W_0 = 17$ | Frame Thickness | $e_f = 1.45 \text{ mm}$ |

**Figure 6:**
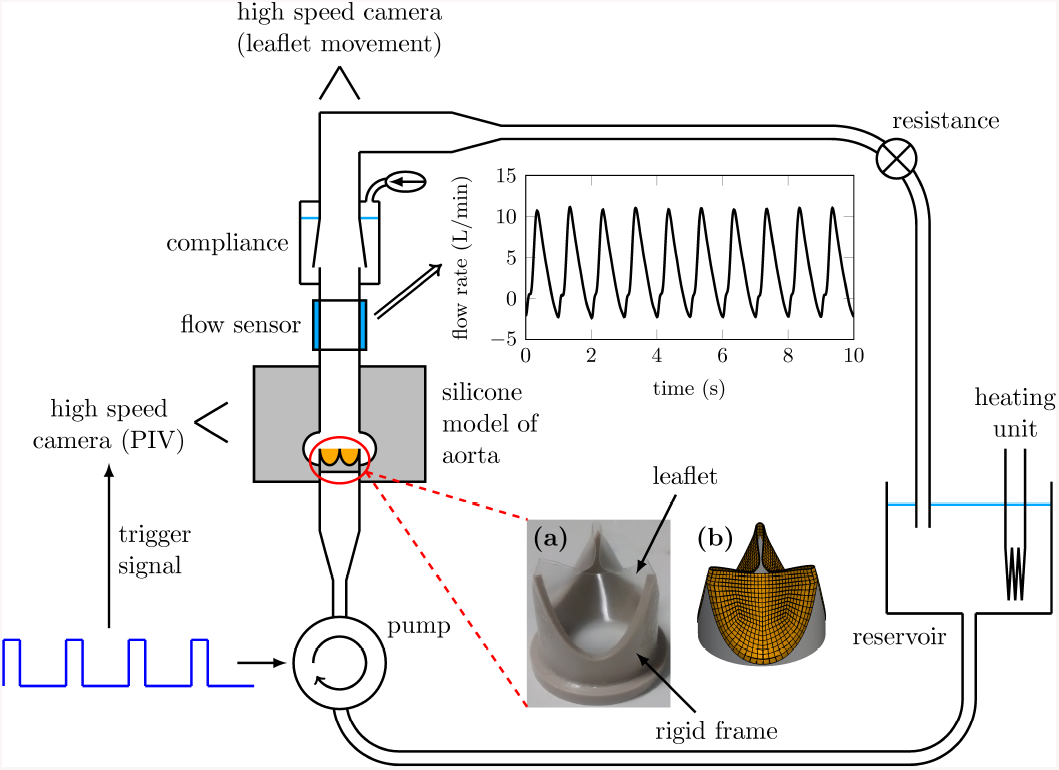
Experimental setup from Sigüenza et al. consisting of a TAVR deployed in a fluid circuit driven by a pulse duplicator, diagram was extracted from Sigüenza et al. [42].

#### 3.2.2 Standard QoIs for BAVR performance

The ISO 5840-3 standard specifies QoIs for evaluation of AVR performance [41]. Among these quantities, the orifice area and TPG are of particular interest in both regulatory and clinical contexts. In this section these quantities are contrasted against experimental data from Sigüenza et al. [42]. While the orifice areas obtained in the present numerical model are lower than the reference experimental data, the maximum TPG is in close agreement with the experimental values and the minimum TPG for the most rigid leaflets is in better agreement than the numerical model from the same work [42]. The effect of leaflet rigidity on these quantities is also analyzed below. Increasing leaflet rigidities are observed to produce smaller mean systolic orifice areas and larger systolic and (absolute) diastolic TPGs, elevating the risk of LVH and cardiac failure.

##### 3.2.2.1 Geometric orifice area

The orifice area is a measure of the degree of opening of the valve leaflets. During systole it is desirable to maximize the orifice area in order to reduce the resistance imposed by the valve during the systolic ejection of blood flow. If systolic orifice area is diminished, then a stronger resistance is imposed on the flow during the ejection phase, loading the left ventricle and elevating the risk of LVH. On the other hand, orifice area must be near zero during diastole to assure that regurgitation is minimized during diastole.

Due to the difficulty of measuring this quantity *in vivo*, there is more than one way of quantifying the orifice area: the effective orifice area (EOA) or the geometric orifice area (GOA), which are respectively calculated from flow quantities or from the geometry itself. The EOA is computed using Gorlin’s formula (53), which can be derived from Bernoulli’s law by neglecting the viscous term, resulting in a function of the mean systolic TPG_sys_, Root Mean Squared (RMS) systolic flow rate *Q*_rms_ and fluid density *ρ*^*f*^ :

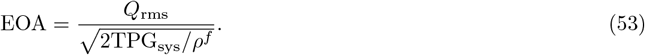

This measure is less accurate than the GOA, but is easier to measure in the clinical context. On the other hand, since the EOA depends on TPGs, it also depends on where pressure is measured on both sides of the valve. Given that the precise location of pressure transducers is unknown, in this work instead the GOA is used to characterize the opening and closing of the leaflets.

###### Algorithm 2 Calculation of geometric orifice area

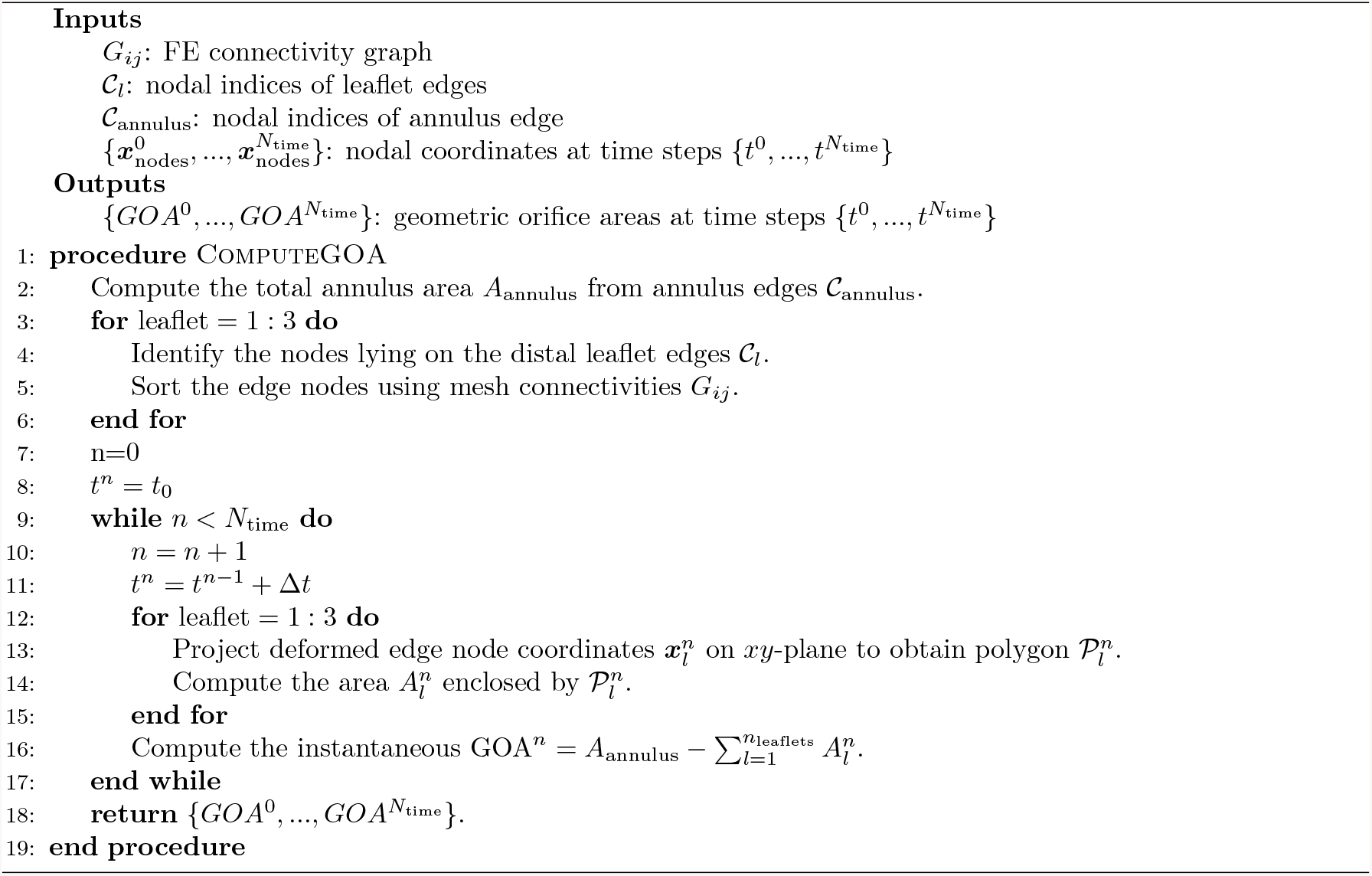

The GOA is defined as the area enclosed by the projection of the leaflet free edges on the aortic cross section. It is not specified how authors compute it in Sigüenza et al. [42], but in this work the instantaneous GOA is computed using Algorithm 2. The algorithm receives as inputs at initial time *t*_0_: the FE nodal connectivity graph *G*_*ij*_, the nodal coordinates 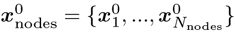 and the indices of surface elements on the leaflet edges 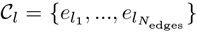. The connectivity graph is defined such that *G*_*ij*_ = 0 if nodes *i* and *j* are disconnected and *G*_*ij*_ = 1 if they are connected, *N*_nodes_ is the total number of nodes, and *N*_edges_ is the number of surface elements on the leaflet edges. For each leaflet *l*, the nodal coordinates of the leaflet edges *𝒞*_*l*_ are projected on the *xy*-plane in order to form the 2D polygon 𝒫_*l*_, whose connectivities are saved once. Also at time *t*_0_, the annulus area *A*_annulus_ is obtained. Then for each time step *t*^*n*^ ∈ [*t*_0_, *t*_*f*_], with time step size Δ*t* and number of time steps *N*_time_, the area 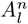 of each polygon 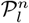 is computed, using the updated nodal coordinates 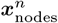 projected on the *xy*-plane. The GOA at each time step is then obtained by summing the areas 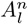 and subtracting them from the annulus area *A*_annulus_. Figure 7 shows snapshots of the valve evolution during the cardiac cycle from the experimental and numerical reference [42] and from this work’s numerical simulations. The shaded leaflets on the plane indicate the areas subtracted at each instant from the total annulus area in Algorithm 2 in order to compute the GOA. The differences in opening and closing dynamics may be attributed to geometry differences between the current geometry and the reference work, snapshots are shown for a qualitative comparison.

**Figure 7:**
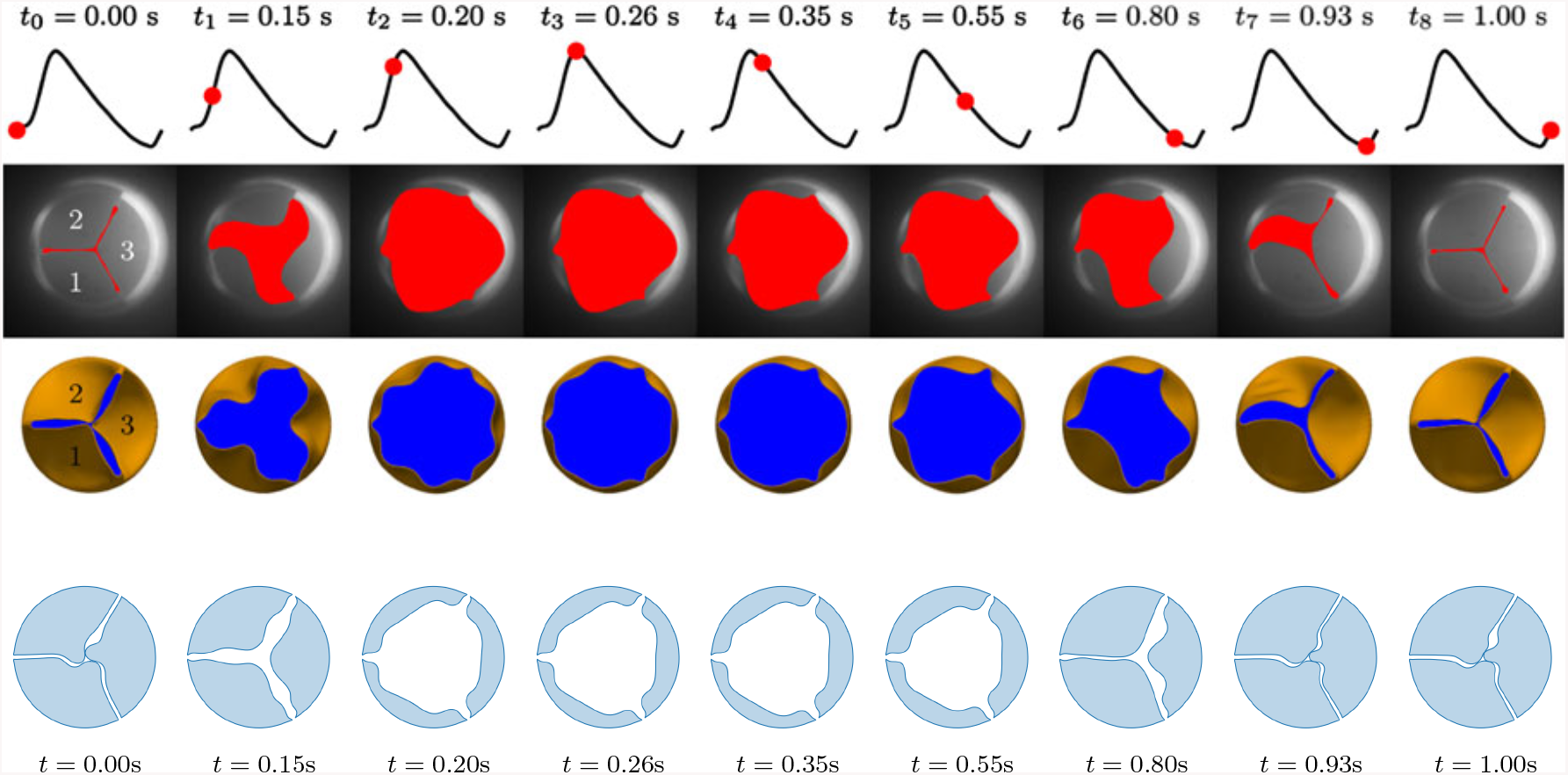
Evolution of valve movement in a single cardiac cycle. From top to bottom: flow rate curve, snapshots from experiments and from numerical simulations in Sigüenza et al. [42], and snapshots from the numerical simulations carried out in this work for the valve with a Young modulus of 1MPa, highlighting leaflet areas used to compute the GOA with Algorithm 2.

Valves of increasing Young moduli were simulated in order to characterize the effect of leaflet rigidity on valve performance. In Figure 8 the valve states are shown both in diastole and systole for the different valve rigidities. In Figure 8b it can qualitatively be observed that during systole the EOA is reduced for more rigid leaflets, reducing systolic function. The GOA is computed to quantify this observation. The instantaneous GOA time series are contrasted in Figure 9a against those provided by Sigüenza et al. [42] for both numerical and experimental data sets. Although the EOA peak values of the present work’s numerical simulations are consistent with reference data, mean values fall below. In addition, the present work shows an earlier valve closure than the reference data. These two differences could be attributed to the differences in valve and aortic chamber geometries, and to the manufacturing process of the experimental valve. In the experiment, a heat treatment was applied to generate the closed valve configuration starting from an open valve initial condition. In the reference numerical simulations a pre-process stage is performed to mimic the experimental initial conditions, but it is not clear if pre-stresses are applied on leaflets as well. If pre-stresses have indeed been applied, this could explain the differences with respect to simulations of the current work. Finally, for more flexible valves in Figure 9a it is evident that leaflet fluttering is more intense than in the reference data. However, as rigidity is increased the fluttering amplitude is decreased.

**Figure 8:**
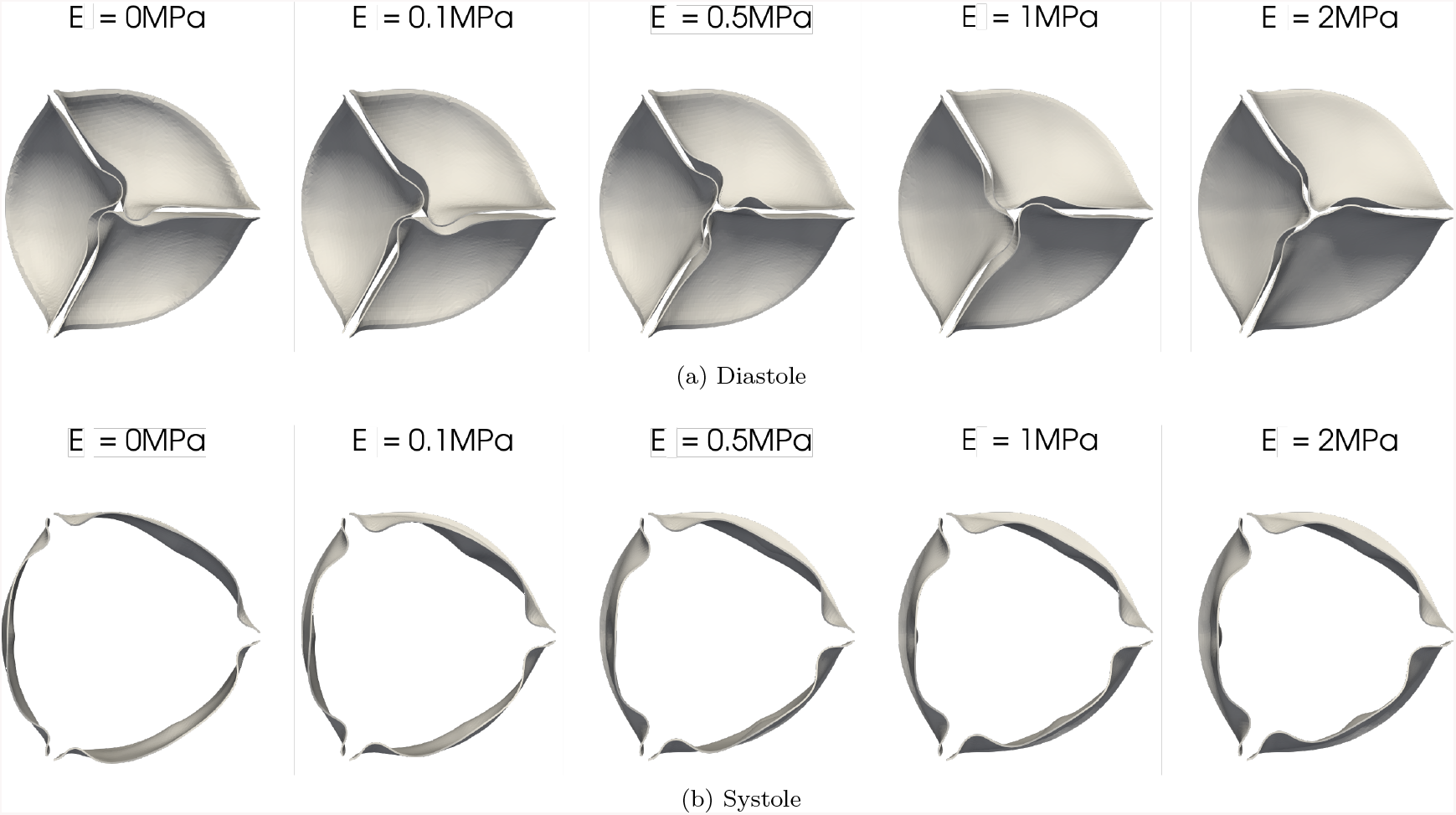
Valve deformation in diastole (top) and systole (bottom) for increasing Young moduli (left to right).

To facilitate the comparison with reference data, the minimum, maximum, time-average over the whole cycle, over the forward flow period (*Q >* 0) and over the backward flow period (*Q <* 0) are extracted for each cardiac cycle. The simulations are run for 3 cardiac cycles. The first cycle is discarded in order to initialize the system. The following 2 cycles are used to compute the phase-averaged statistics shown in Figure 9b. This figure shows that for the range of Young moduli studied, the minimum and maximum GOAs intersect the experimental values, showing consistency with the reference data. Moreover, all valves simulated in the current work comply with the minimum EOA established by the ISO 5840-3 standard [41], *ie:* EOA_*min*_ = 1.45cm^2^ for 25mm valve diameters.

**Figure 9:**
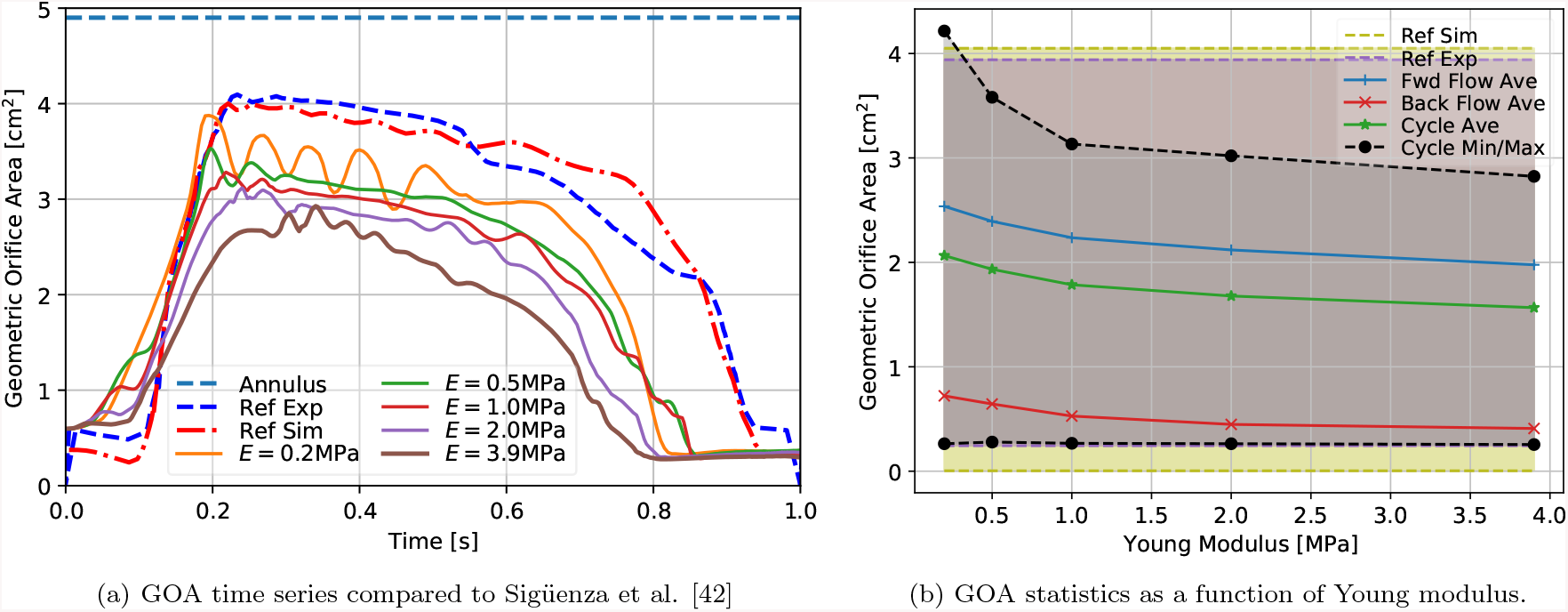
GOA for different Young moduli of valve leaflets for both (a) time series (continuous lines) compared to reference values from Sigüenza et al. [42] (dashed for experimental and point-dashed for numerical data) with the pointed line indicating the fixed annulus area, and (b) phase-averaged over the forward-flow period (blue +’s), backflow period (red x’s), entire cycle (green *’s) and the cycle minimum and maximum (black dashed for this work’s numerical data, pink and violet for the experimental and numerical references [42]). Pink and violet shading are used to show range of values for experimental and numerical data respectively.

Regarding the effect of rigidity on valve opening, in Figure 9b it is observed that with increasing rigidity the maximum GOA (during systole) is significantly decreased while the minimum GOA (during the backflow period) remains approximately constant with a slight decrease. The former result agrees with the well documented observation that calcified leaflets (ie: more rigid) increase the risk of diminished systolic EOAs and intravalvular regurgitation [66]. Moreover, for applications in tissue engineered heart valves, this curve may be generated for a given valve model to determine the range of possible Young moduli which will yield the desired range of systolic orifice areas.

##### 3.2.2.2 Transvalvular pressure gradient

In this section, TPGs are computed by averaging the fluid pressure at cross-section slices 2cm upstream (*P*_up_(*t*^*n*^)) and 2cm downstream (*P*_down_(*t*^*n*^)) of the valve plane, and computing the difference between them at each time step *t*^*n*^: TPG(*t*^*n*^) = *P*_*up*_(*t*^*n*^) − *P*_*down*_(*t*^*n*^). The valve plane is the plane formed by the free edges of the leaflets when closed. Phase-averaged statics from this time series are extracted, as for the GOA in the previous section. The resultant TPG statistics are visualized in Figure 10 as a function of the leaflets’ Young moduli. At this point, other epistemic errors may arise when comparing to the reference experiment since the exact location of pressure transducers are not given in Sigüenza et al. [42]. Nonetheless, as can be seen in Table 3, considering the range of Young moduli evaluated, the resultant TPG_max_ intersects the numerical reference value (4.9mmHg) and is in close agreement to the experimental value (5.28mmHg). For TPG_min_, the present numerical model is closer to the experimental value than the numerical model from Sigüenza et al. [42]. Nonetheless, it is hypothesized that the difference in TPG_min_ between both numerical models and the experimental data is originated by the fact that none consider the compliance and resistance models at the aortic outlet.

**Table 3:** Minimum and maximum GOA and TPG, from top to bottom: this work’s numerical simulations of increasing leaflet rigidity, numerical simulations and experimental results from Sigüenza et al. [42].

| Source | $GOA_{min}$ [cm<sup>2</sup>] | $GOA_{max}$ [cm<sup>2</sup>] | $TPG_{min}$ [mmHg] | $TPG_{max}$ [mmHg] |
| --- | --- | --- | --- | --- |
| $E = 0.2$ MPa | 0.263 | 4.22 | -8.80 | 4.93 |
| $E = 0.5$ MPa | 0.279 | 3.58 | -18.0 | 5.02 |
| $E = 1.0$ MPa | 0.267 | 3.13 | -25.7 | 4.95 |
| $E = 2.0$ MPa | 0.263 | 3.02 | -38.0 | 6.11 |
| $E = 3.9$ MPa | 0.255 | 2.82 | -51.9 | 5.30 |
| Sigüenza Simulation | 0.244 | 4.05 | -21.05 | 4.9 |
| Sigüenza Experiment | 4.04e-3 | 3.94 | -93.35 | 5.28 |

**Figure 10:**
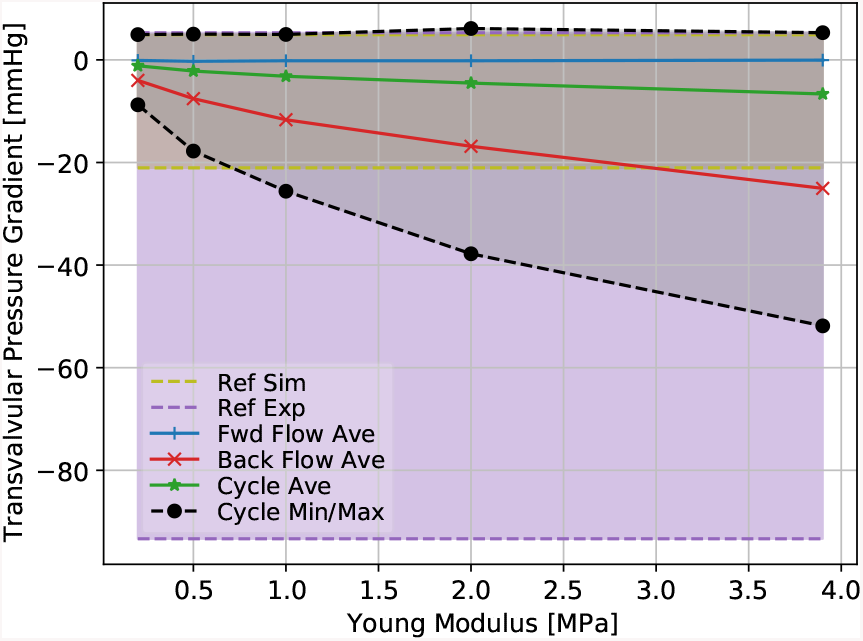
Phase-averaged TPGs as a function of the Young moduli of valve leaflets and comparison to reference experimental and numerical values [42].

It is observed in Figure 10 that the maximum TPG grows with the Young modulus, which indicates a higher resistance to the flow. On the other hand, the minimum TPG increases in modulus as the Young modulus is increased, which indicates that more rigid leaflets are able to withstand higher pressure gradients during the backflow period, a desired characteristic for valve prostheses. This shows that although softer leaflets provide larger systolic EOA and lower systolic TPGs, a moderate degree of rigidity is desirable in order to maintain the negative TPGs during the backflow period. In the following section an analysis is carried out on the effect of leaflet rigidity on biomarkers associated to thrombogenic risk.

#### 3.2.3 Thrombosis QoIs

In this section the effect of leaflet rigidity on two QoIs associated to thrombogenic risk are studied, namely residence time and shear rate. It is observed that more rigid valves intensify stagnation points in diastole and peak shear rate regions in both diastole and systole, resulting in a higher risk of thrombus formation. This provides a framework for directly testing the effect of different BAVR model parameters on the risk of thrombus formation, a subject of yet unclear clinical consequences [7, 8].

##### 3.2.3.1 Residence time

The residence time (*T*_*R*_) of a fluid parcel is the total time that the parcel has spent inside a control volume. It therefore quantifies the degree of washout of old fluid at each point in space. Although the increased *T*_*R*_ does not lead to thrombus formation by itself, it is known that once the clotting process is triggered, for instance by blood contact with the foreign surfaces of TAVRs, thrombosis is more likely to occur in low flow regions characterized by large *T*_*R*_’s [17, 67, 68]. In this work *T*_*R*_ is computed using a Lagrangian approach: a passive scalar is advected by the flow using equation (54),

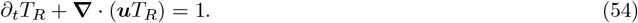

Dirichlet boundary conditions *T*_*R*_ = 0 are set at the inlet, while zero-flux Neumann boundary conditions, ***n*·∇***T*_*R*_ = 0, are set at the outlet and vessel walls (*ie*: adiabatic boundary conditions).

Thrombus formation is a process which is initiated by nucleation [69], therefore it is important to consider local activation points to determine the thrombogenic risk. Although in Figure 12 volume and phase-averaged residence times are slightly reduced for more rigid leaflets, peak residence time regions in Figure 11 are observed to intensify with leaflet rigidity, particularly in the sinus of Valsalva (SoV) region, reducing blood washout. Peak residence time regions are identified as regions with *T*_*R*_ *>* 1.5*T*_cycle_, where *T*_cycle_ = 1s. In order to fully comprehend the thrombogenic risk, the effect of shear rate must be taken into account as well. This quantity is analyzed in the next section.

**Figure 11:**
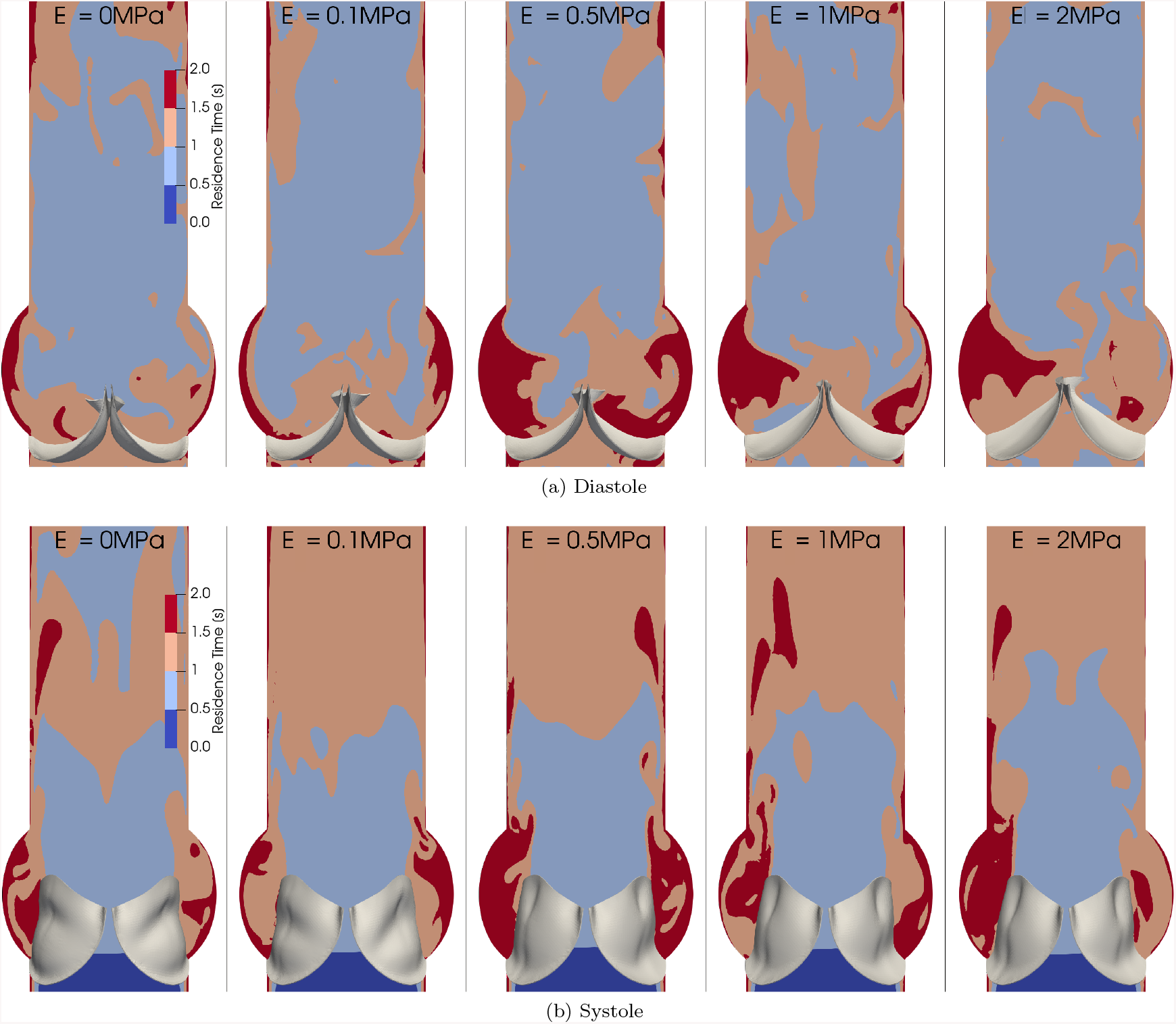
Residence time for increasing Young moduli (left to right) in diastole (top) and systole (bottom).

**Figure 12:**
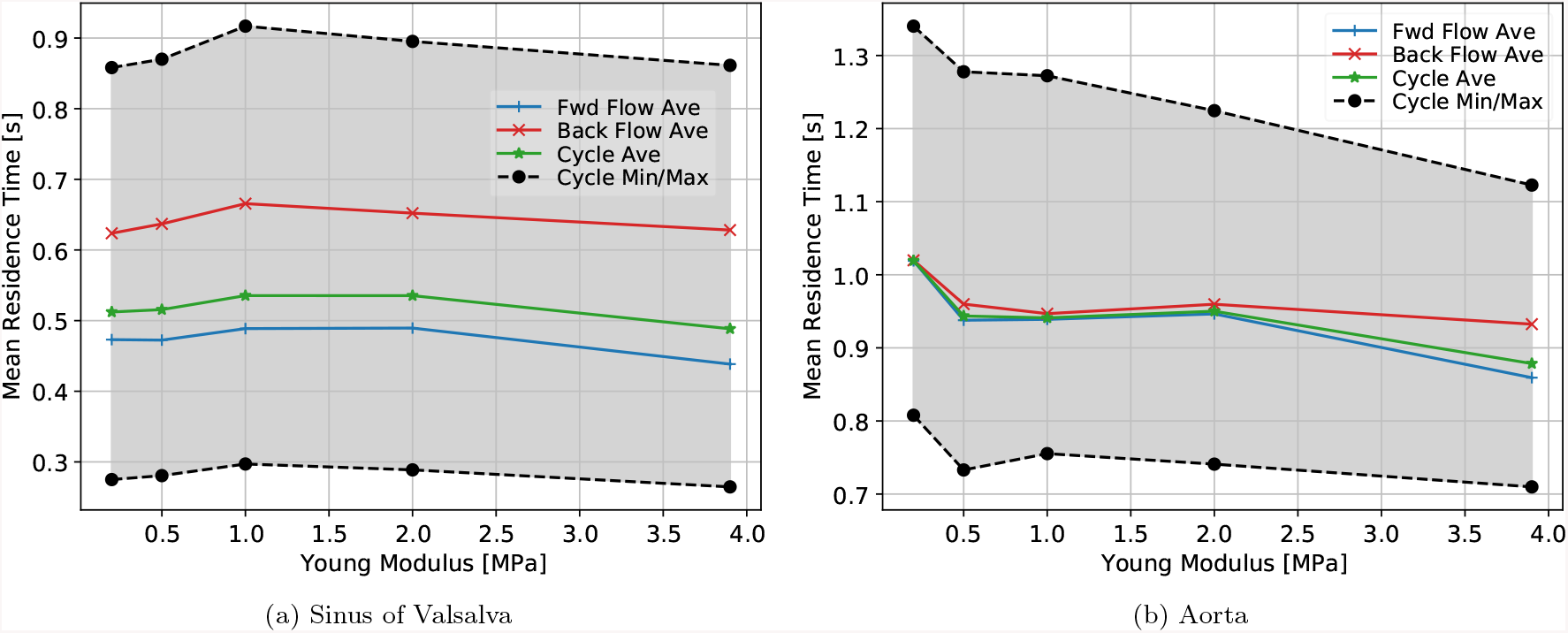
Volume- and phase-averaged residence time as a function of the Young moduli of valve leaflets.

##### 3.2.3.2 Shear rate

As described in the introduction, thrombi are usually formed in regions of stagnation, that is long residence times, and low or high shear. Shear rate has been associated to thrombogenic risk [20]. It is a convenient measure since it is a Galilean invariant, which implies that it is not modified by a linear rigid body motion. It is computed as the squared root of the second invariant of the symmetric strain tensor *Q*_*S*_:

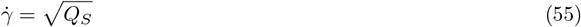

where 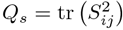 is the second invariant and *S*_*ij*_ = *∂*_*i*_*u*_*j*_ + *∂*_*j*_*u*_*i*_ is the symmetric strain tensor. Figure 14 shows the effect of the Young modulus on the volume- and phase-averaged shear rate in the sinus of Valsalva and aorta.

**Figure 13:**
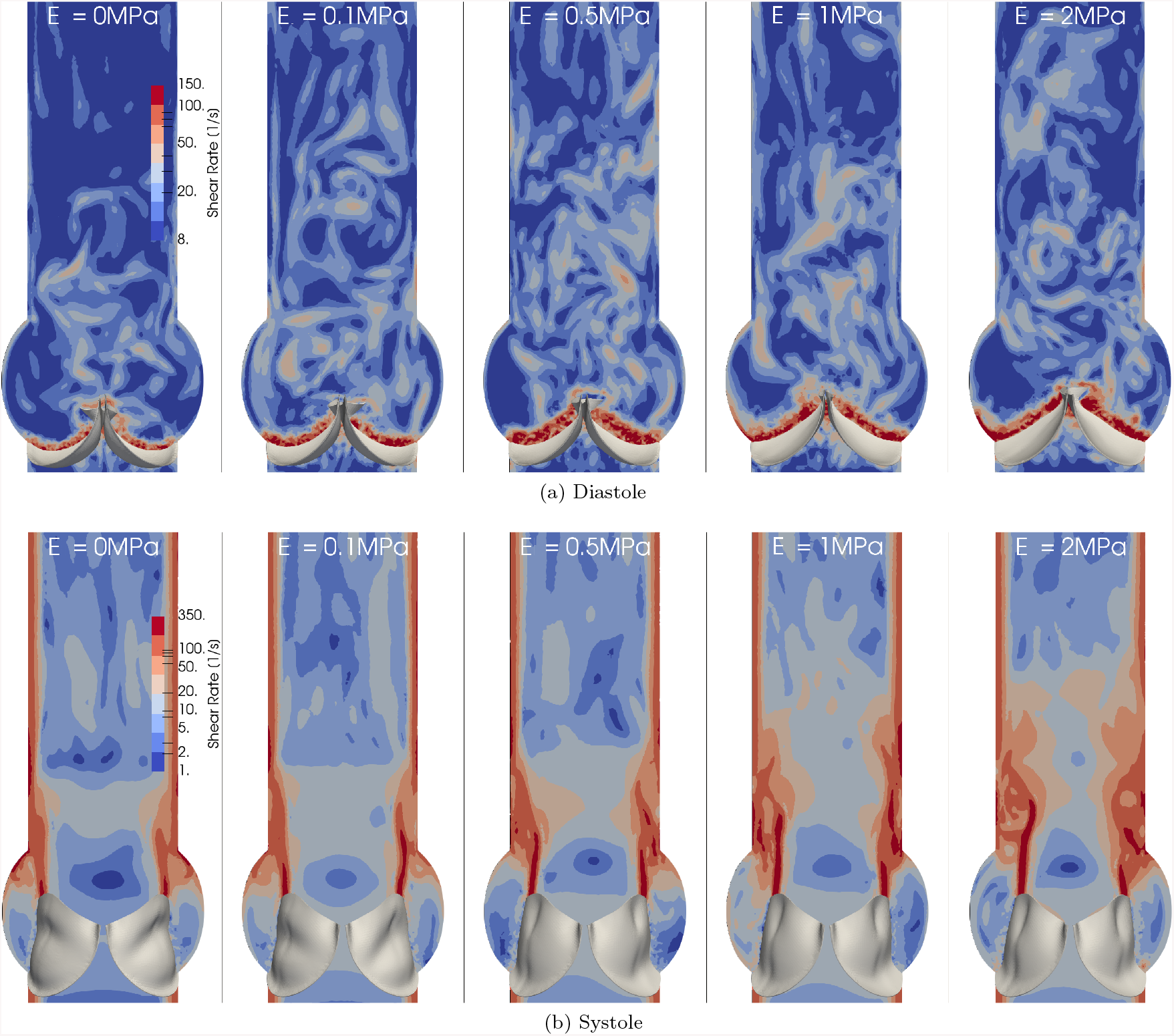
Shear rate for increasing Young moduli (left to right) in diastole (top) and systole (bottom).

**Figure 14:**
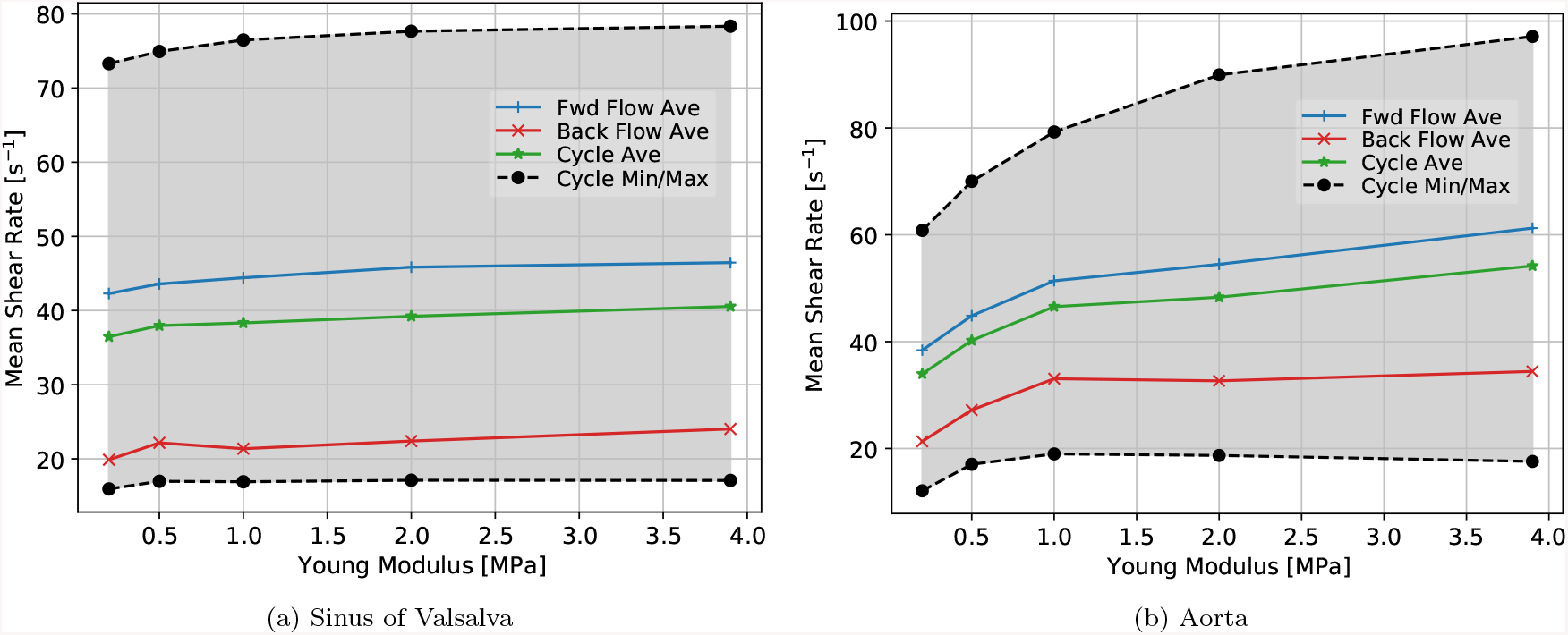
Volume- and phase-averaged shear rate as a function of the Young moduli of valve leaflets.

As mentioned in the introduction, red and white clots are formed at low (*<* 50s^−1^) and high (*>* 5000s^−1^) shear rates respectively [12]. Figure 14 shows the volume- and phase-averaged shear rate in the sinus of Valsalva and aorta as a function of the leaflet Young modulus. In these cases average shear rates are within 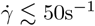, that is, within the range of shear rates associated to red clot formation. Here shear rate increases with the Young modulus, which can be explained by more rigid leaflets increasing the degree of flow obstruction, while forming more vortical structures in a moderately turbulent regime. Moreover, when visually inspecting the shear rate fields in Figure 13, extreme values are intensified with leaflet rigidity. Recalling that in the previous section (3.2.3.1) peak residence time regions are also intensified with rigidity, it can be concluded from this analysis that in the current working conditions, an increase in leaflet rigidity produces an increase in risk of blood clot formation according to the considered biomarkers. Although at lower Reynolds numbers, a similar effect of leaflet rigidity on thrombogenic risk has been observed in an *in vitro* experiment for venous valves [70]. Further *in vitro* and *in vivo* studies should be carried out to confirm this observation for native and bioprosthetic aortic valves.

#### 3.2.4 Calcification biomarker: von Mises Stresses

As described in the introduction, mechanical stress has been extensively associated to valve calcification and failure. In this section the von Mises stresses in leaflets are computed to assess the risk of valve calcification, they are computed as

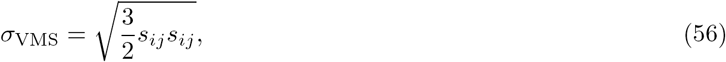

where *s*_*ij*_ is the deviatoric stress tensor of the structure. In Figure 15 the volume- and phase-averaged von Mises stresses in the leaflets are shown as a function of the Young modulus of the material. It is observed that stresses grow with the Young modulus as expected given that more rigid materials yield higher stresses for the same strains. Furthermore, when inspecting the von Mises stress fields visually in Figure 16, it is notable that higher stresses become more concentrated on the leaflet comissures and attachments. These stress hot-points are hypothesized to become points of tissue wear-out, or catalyzers of the calcification process.

**Figure 15:**
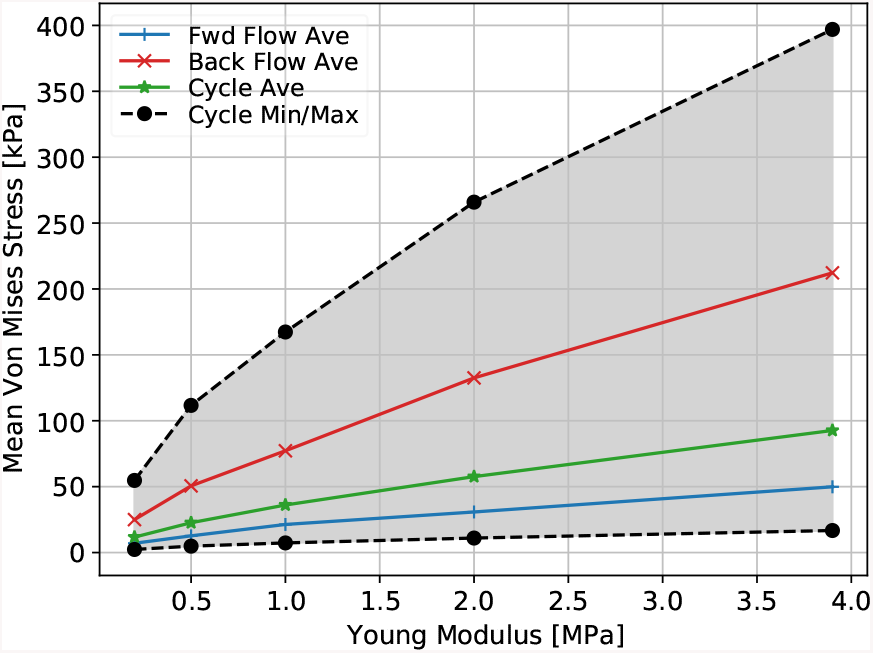
Mean leaflet von Mises stresses as a function of the Young modulus of valve leaflets.

**Figure 16:**
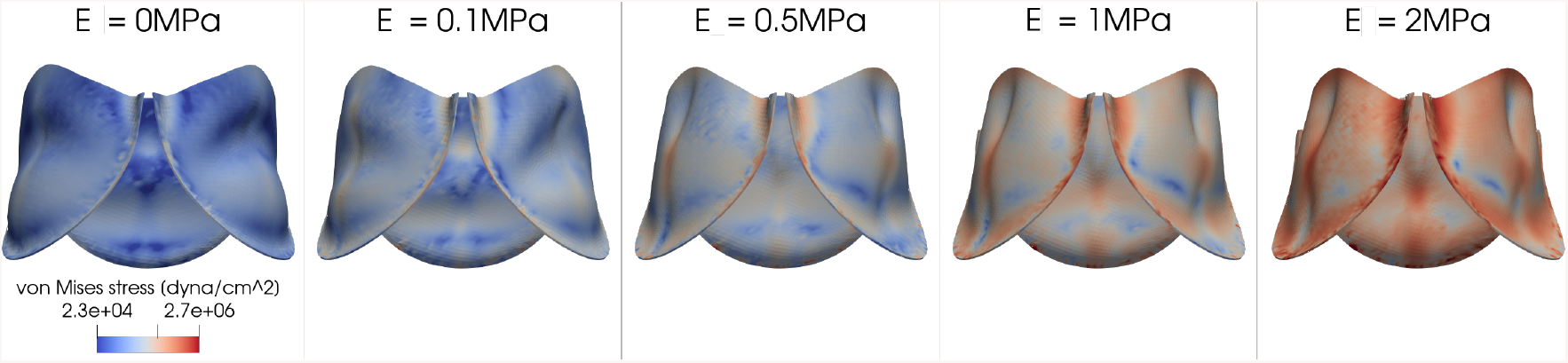
von Mises stresses in systole for increasing Young moduli (left to right).

### 3.3 Effect of eccentricity on aortic valve replacement

This section presents a study on the effect of the eccentricity of the aortic root on the QoIs studied so far, related to valve prosthesis performance, and to thrombogenic and calcification risks. Results are contrasted against the experimental *in vitro* study by Kütting et al. [32].

#### 3.3.1 Setup

In the experimental work by Kütting et al. [32], authors deploy 2 transcatheter heart valves (Medtronic CoreValves) of diameters 26mm and 29mm in silicone compartments of diameters 23mm and 26mm respectively, that is, within the ranges of oversize indicated by the manufacturer. The silicone compartments emulate the aortic root geometry. They were designed with three different eccentricity indexes (EIs) of the aortic annulus, while maintaining the area constant: *EI* = 0 (circular), 0.25 (moderate eccentricity) and 0.33 (highly eccentric). The eccentricity index is computed as *EI* = 1 − *D*_*min*_*/D*_*max*_, with *D*_*min*_ and *D*_*max*_ the elliptic short and long axes respectively. The measurements were performed with water in a pulse duplicator using three different physiological flow rate waveforms corresponding to low cardiac output (CO = 3lmin^−1^ at 120 BPM), physiological flow (CO = 5lmin^−1^ at 70 BPM) and physical exercise conditions (CO = 8lmin^−1^ at 110 BPM). The pulse duplicator was designed to model the flow conditions of the left heart. It is equipped with a passively filling left atrium, mitral valve, silicone ventricle, interchangeable aortic roots and adjustable peripheral impedance as shown in Figure 17. The stroke volume of the ventricle and the beat rate can be adjusted to alter the flow rate of the system. Following the guidelines of the ISO 5840-3 standard [41], QoIs reported include: regurgitation flow, systolic EOA and systolic TPG. The current work replicates the settings of the *in vitro* study for the 23mm diameter aortic root (26mm TAVR), with some simplifications:

**Figure 17:**
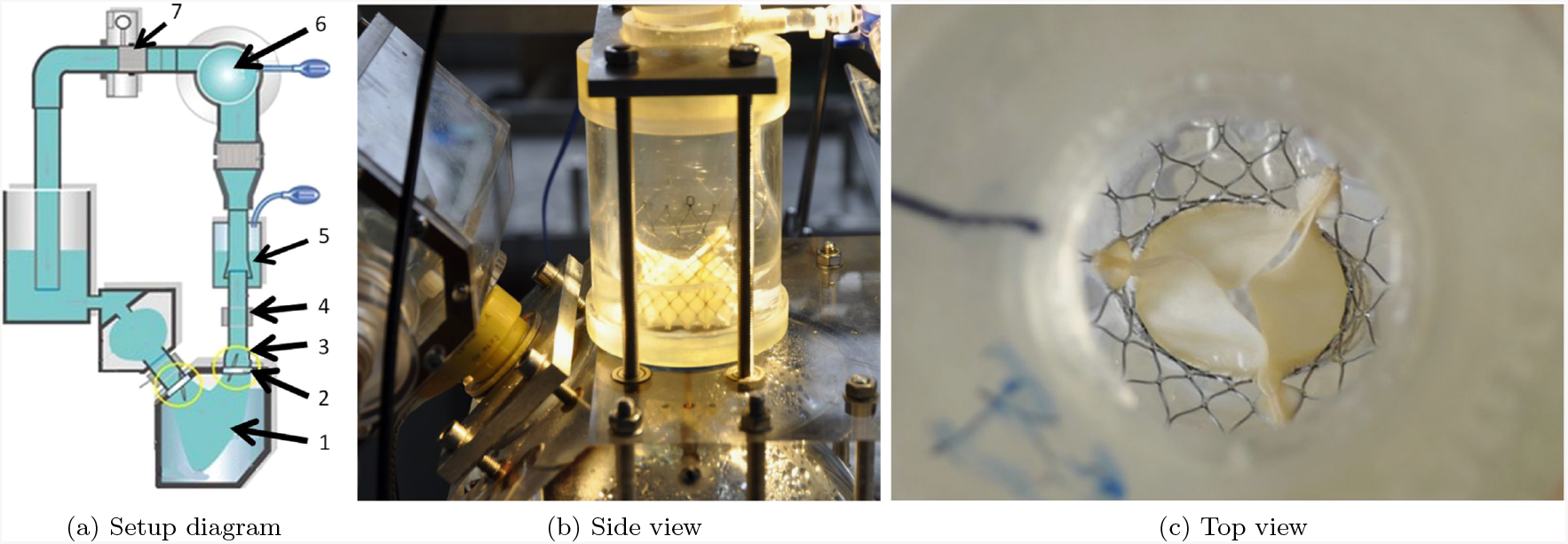
Experimental setup from Kütting et al. [32]

1. the aortic root is considered to be rigid,
2. the geometry of the reference work is approximated from the available information, such as the aortic annulus nominal diameters and eccentricities which are respected,
3. flow rate is imposed at the inlet on the left ventricle side and a constant pressure at the outlet of the aortic root, while the reference *in vitro* study imposes compliant pressure boundary conditions at the outlet which simulates the compliance of the systemic circulation,
4. valve leaflets are modeled using an isotropic Neohookean material model,
5. no left ventricle is included in the model,
6. the outflow is modeled as an open boundary instead of compliant, and
7. the eccentricity is modeled by scaling the valve prostheses asymmetrically

The effect of the aortic root eccentricity on the following QoIs associated to TAVR performance is analyzed: TPG and GOA for different cardiac outputs (COs). In Kütting et al., the orifice area is quantified using Gorlin’s formula (53) to compute the EOA [32]. As explained in section 3.2.2.1, since the EOA depends on the location of the pressure transducers, which are not specified in the reference work, here the geometric measure of the orifice area is used instead, which is computed once again using Algorithm 2. Given that flow rate is imposed at the inlet and that the fluid is incompressible, the flow rate at each cross section in the domain is trivially determined, and is the same at each cross section at any given point in time. Therefore, regurgitation flow (also known as regurgitant volume), which is computed as the volume of backward flow across the aortic valve during the cardiac cycle, is also trivially determined by the prescribed boundary conditions. To compute regurgitation flow without prescribing it a priori, pressure boundary conditions should be used instead at inlet and outlet. Since these boundary conditions are out of the scope of the current work, regurgitation flow is excluded from this study. As in section 3.2, the reference *in vitro* study is extended, to analyze the effect of eccentricity on the thrombogenic and calcification biomarkers analyzed previously, that is, residence time, shear rate and von Mises stresses.

#### 3.3.2 Standard QoIs for BAVR performance

This section analyzes the effect of eccentricity on the QoIs determined by the ISO 5840-3 for valve performance assessment, that is, orifice area and TPG. With increasing eccentricity, it is observed that GOA decreases in healthy conditions while TPG increases in all cases, which indicates a decrease in valvular function for more eccentric annuli.

##### 3.3.2.1 Geometric orifice area

The GOA is computed using Algorithm 2 as in section 3.2.2.1 at each instant for the valves shown in Figure 18. The phase-averaged values are then compared to the reference values in Figure 19. It is observed that tendencies of decreasing forward flow orifice areas with increasing eccentricity are reproduced in the healthy rest and exercise working conditions, that is, *CO* = 5 and 8lmin^−1^ respectively (see purple and blue curves in Figures 19b-c). The positive correlation between eccentricity and orifice area observed in the pathological flow rate case (*CO* = 3lmin^−1^) is not commented upon in the experimental reference [32], but it is not evident why this tendency is not reproduced in the current numerical simulations. For the healthy working conditions, an offset is observed between the numerical and experimental curves, which could be attributed to systematic differences between geometries. Moreover, the absence of pre-stress is another factor which may be elevating the measured GOA in the numerical results. However, these results confirm that in healthy cardiac output conditions an increase in eccentricity index decreases the orifice area, worsening valve performance.

**Figure 18:**
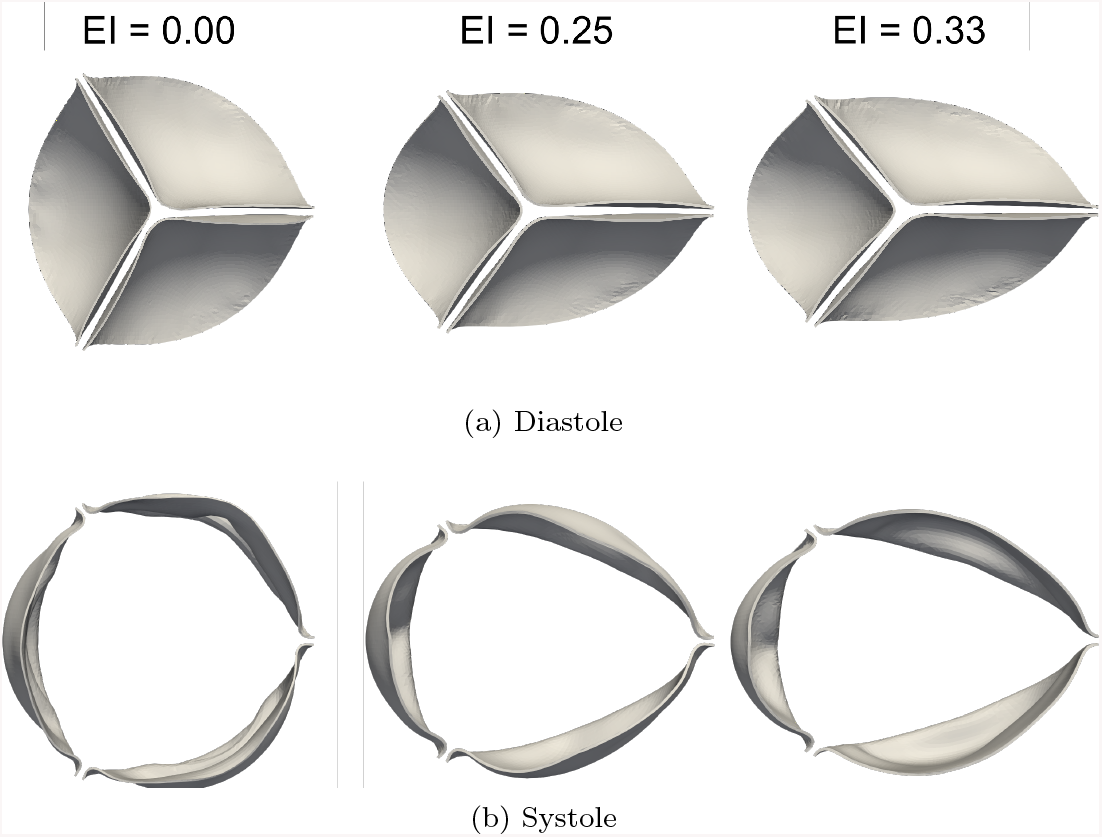
Valve deformation in (a) diastole, and (b) systole, for increasing eccentricities (left to right) and for conditions corresponding to a healthy patient at rest: *CO* = 5lmin^−1^.

**Figure 19:**
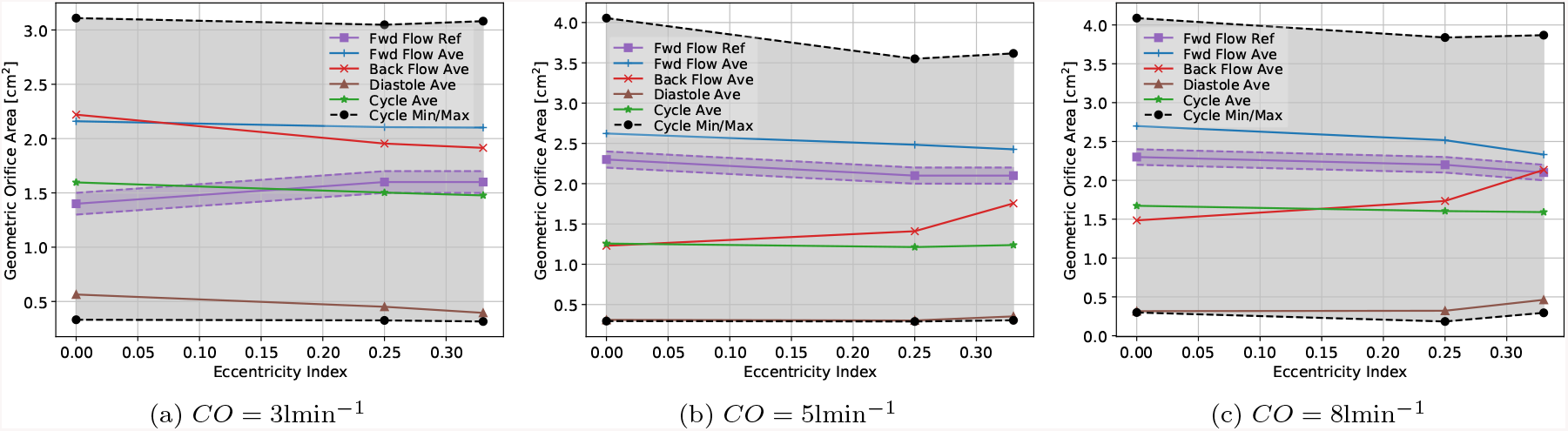
Phase-averaged GOA as a function of the eccentricity index and compared to EOA from the *in vitro* experiment [32] in different flow rate conditions corresponding to cardiac failure (*CO* = 3lmin^−1^), healthy patient at rest (*CO* = 5lmin^−1^) and healthy patient in exercise conditions (*CO* = 8lmin^−1^).

##### 3.3.2.2 Transvalvular pressure gradient

TPG is computed as in section 3.2.2.2 for each cardiac output and each eccentricity index. The results are observed in Figure 20. Since the geometry of the experimental setup also includes the left ventricle, and the locations of the experimental pressure transducers are unknown, absolute values of pressure are not directly comparable to the experimental reference. Instead, comparisons are carried out between the mean systolic TPGs normalized by the value corresponding to the circular case (TPG_circular_).

**Figure 20:**
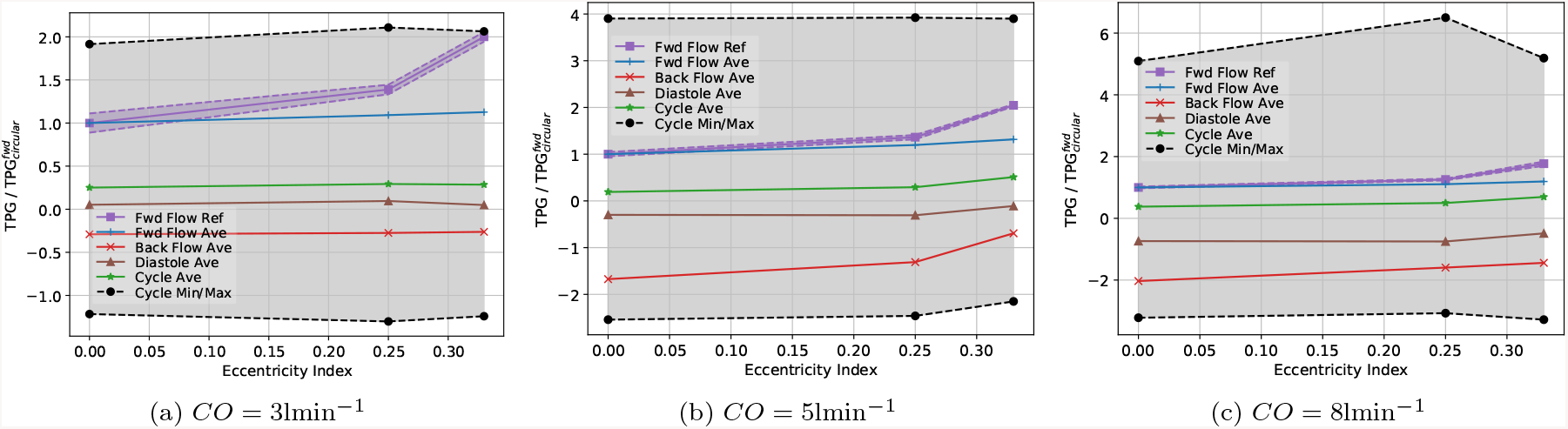
Normalized phase-averaged TPG as a function of the eccentricity index and compared to EOA from the *in vitro* experiment [32]. Values are normalized by the corresponding value in the circular case to clarify comparison. The increasing tendency of systolic TPG with eccentricity is reproduced in the numerical experiments for the healthy rest and exercise conditions, *ie: CO* = 5 and 8lmin^−1^.

The tendency of increasing mean forward flow TPG for increasing eccentricity is reproduced in the normalized numerical data presented in Figure 20. A good agreement is particularly observed for the 23mm diameter cases with 5 and 8lmin^−1^ cardiac outputs. This tendency indicates that for healthy patient conditions both at rest and in exercise, an increase in eccentricity of the annulus is translated into a larger TPG during systole. The larger systolic TPG due to a narrowing of the aortic valve orifice increases the workload on the left ventricle. In response to increased after-load, LVH can develop to maintain cardiac function. The LVH response ultimately decompensates with progressive cell death and fibrosis, driving the transition to symptoms, heart failure, and adverse cardiovascular events [71, 72, 73]. As a consequence, the present results indicate that more eccentric annulus could enhance the risk of LVH.

#### 3.3.3 Thrombosis QoIs

In this section, it is observed that increasing eccentricity enhances peak residence time and shear rate regions, thus increasing the thrombogenic risk according to these biomarkers of leaflet thrombosis.

##### 3.3.3.1 Residence time

As explained in section 3.2.3.1, thrombosis is more likely to occur in low flow regions characterized by large residence times [17, 67, 68]. Figure 22 shows the time- and volume-averaged residence time in the sinus of Valsalva for different cardiac outputs. Here residence time reaches a minimum for the intermediate eccentricity, *EI* = 0.25. However, when visually inspecting the snapshots of residence time at diastole and systole in Figure 21, it is evident that in both diastole and systole, regions of peak residence time are systematically intensified as valves become more eccentric. Similarly as in section 3.2.3.1, given that thrombus formation is triggered from nucleation points, it is important to note that while mean values do not show a monotonic tendency for residence time as a function of eccentricity, peak residence time does show a monotonically increasing tendency from the circular to the most eccentric configuration. This result suggests that a high degree of eccentricity not only affects the valve performance, but also the washout in the sinus of Valsalva.

**Figure 21:**
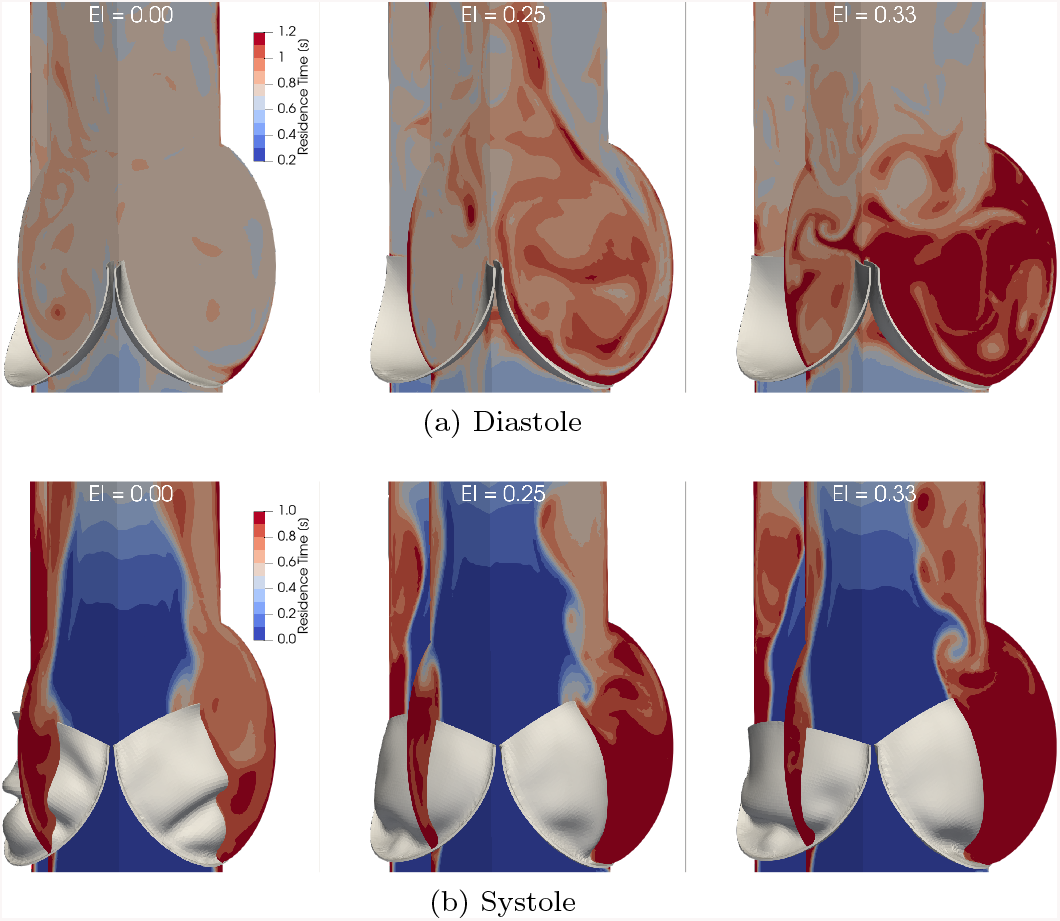
Residence time in (a) diastole, and (b) systole, for increasing eccentricities (left to right) and for conditions corresponding to a healthy patient at rest: *CO* = 5lmin^−1^.

**Figure 22:**
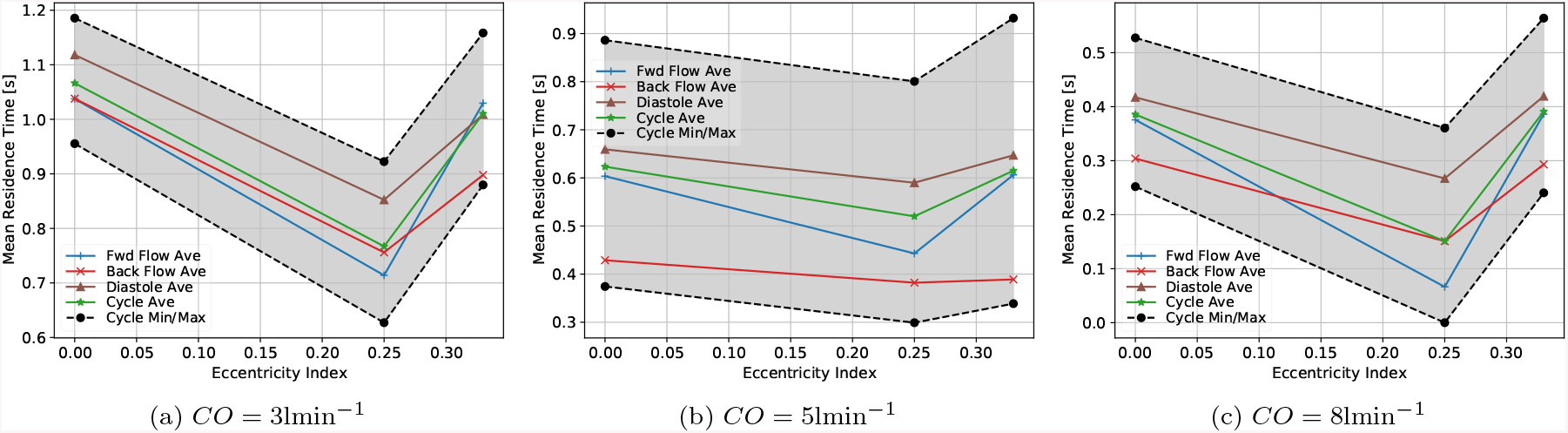
Volume- and phase-averaged residence time in the sinus of Valsalva as a function of the eccentricity index.

##### 3.3.3.2 Shear rate

Shear rate is associated to thrombogenic risk [20] due its role in platelet activation. Figure 23 shows snapshots of the shear rate in diastole and systole for the *CO* = 5lmin^−1^ flow rate conditions. Here it is observed that as eccentricity increases, the number of vortical structures transitioning towards turbulence are also increased, generating more shear rate in the flow. Nonetheless, shear rate levels are maintained below 800s^−1^, thus still attributed to red clot formation. In diastole peak shear rate structures are located on the aortic faces of leaflets, in the sinus of Valsalva and downstream in the aorta. The sinus of Valsalva and aortic faces of the leaflet also present long residence times for the most eccentric valves in Figure 21a. These regions are therefore considered of high thrombogenic risk. This hypotheses is confirmed when analyzing the predominant thrombus formation risk points in TAVIs (see Demarchena et al. [74]), which are seen to form on the aortic leaflet faces. In systole, the largest shear rates are located on the leaflet trailing edges at the sino-tubular junction, where vortices are formed. In Figure 21b, it is notable that the sino-tubular region also presents long residence times, providing another potential high thrombogenic risk hot-point.

**Figure 23:**
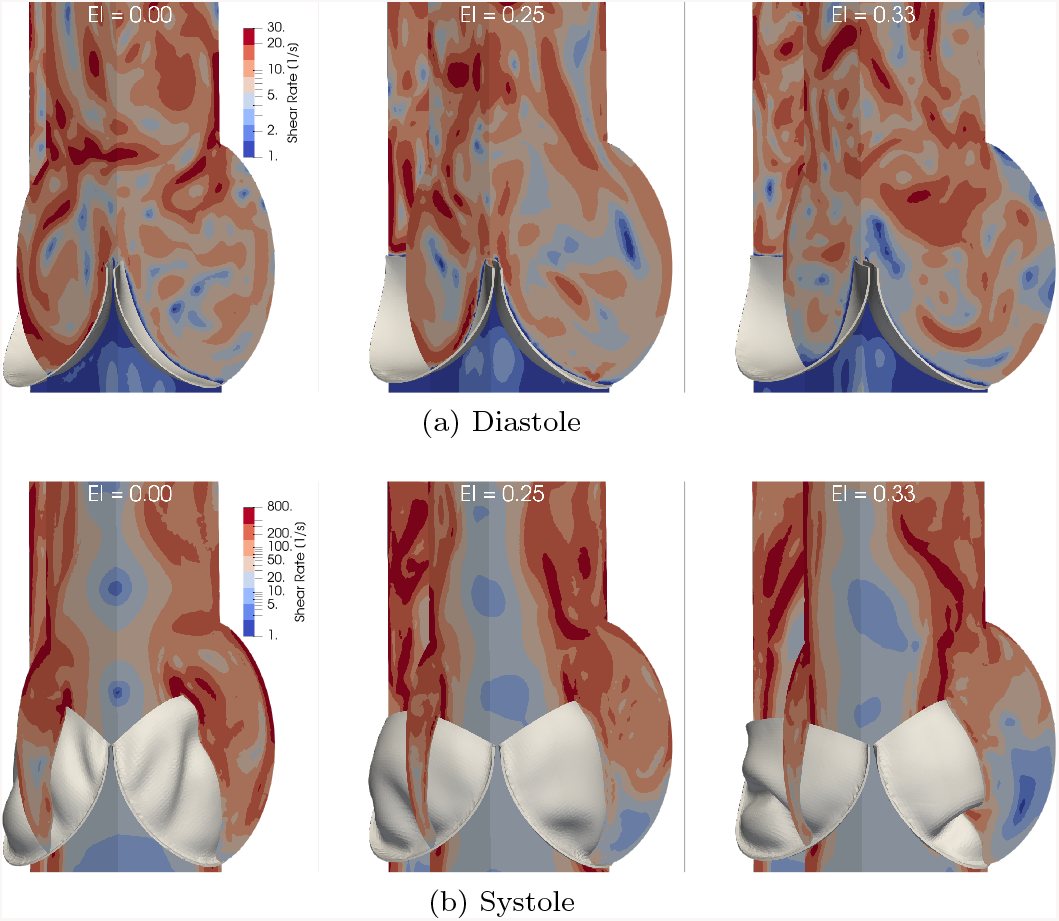
Shear rate in (a) diastole, and (b) systole, for increasing eccentricities (left to right) and for conditions corresponding to a healthy patient at rest: *CO* = 5lmin^−1^.

In Figure 24, time- and volume-averaged residence time in the sinus of Valsalva and aorta are presented for different cardiac outputs. It is interesting to observe that tendencies of shear rate with respect to eccentricity are inverted between the sinus of Valsalva and the aorta. For all cardiac outputs, in the sinus of Valsalva the intermediate eccentricity *EI* = 0.25 yields the maximum shear rate, while for the aorta this same eccentricity produces the minimum shear rate. Therefore, a moderate degree of eccentricity could maximize platelet activation rate in the sinus of Valsalva while minimizing it in the aorta.

**Figure 24:**
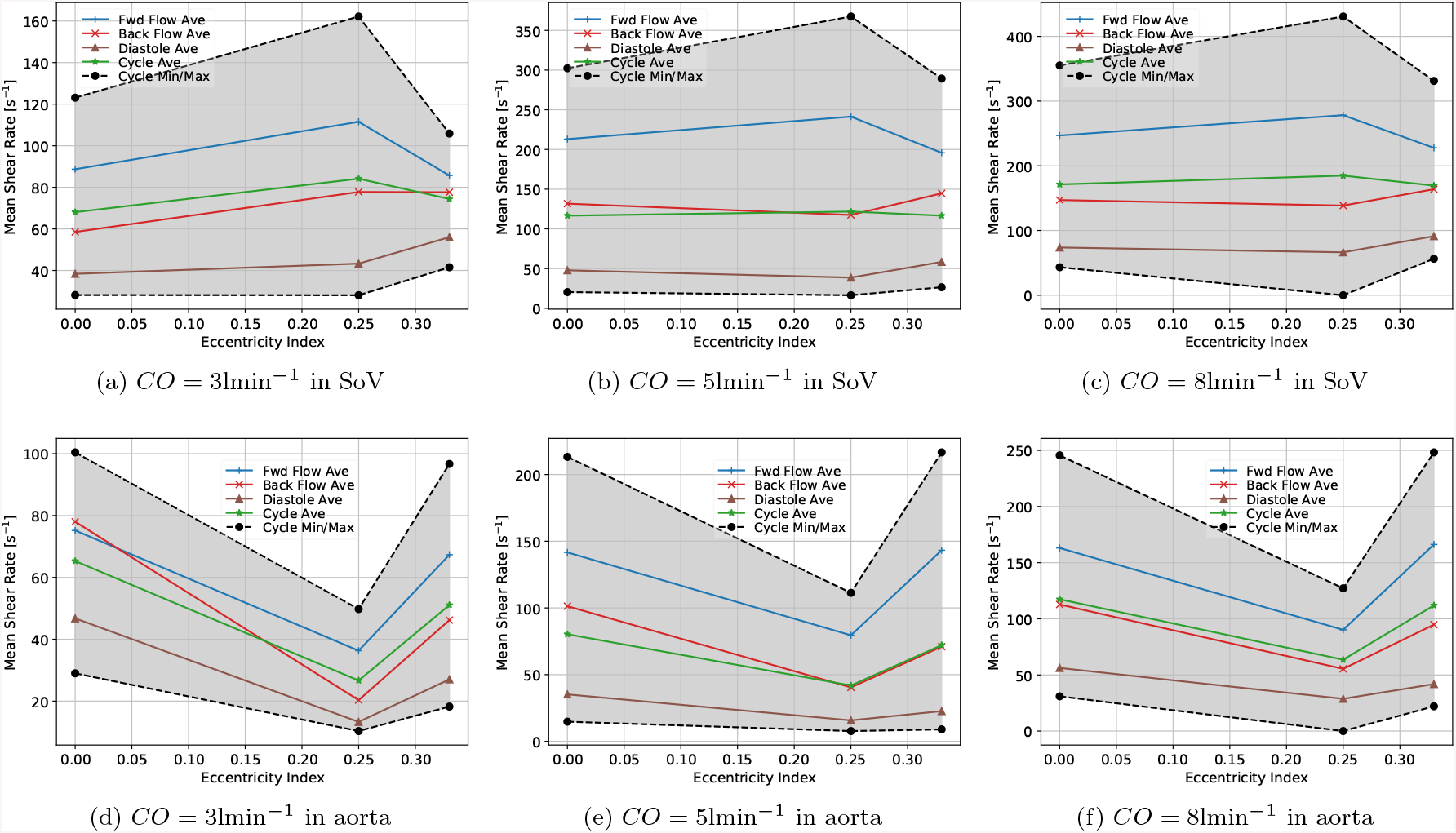
Volume- and phase-averaged shear rate as a function of the eccentricity index of the aortic annulus in the sinus of Valsalva (above) and aorta (below) for different cardiac outputs (increasing left to right).

#### 3.3.4 Calcification biomarker: von Mises Stresses

As explained in 3.2.4, leaflet stresses have been related to prostheses calcification and structural failure. The von Mises stress is used to quantify the degree of stress affecting the leaflets as a function of the eccentricity index of the aortic root. In Figure 26, it can be observed that for the lowest (pathological) cardiac output, an increase in the eccentricity of the aortic root is translated to an increase in the von Mises stresses. This tendency is inverted in the healthy rest and exercise working conditions, that is *CO* = 5 and 8lmin^−1^.

**Figure 25:**
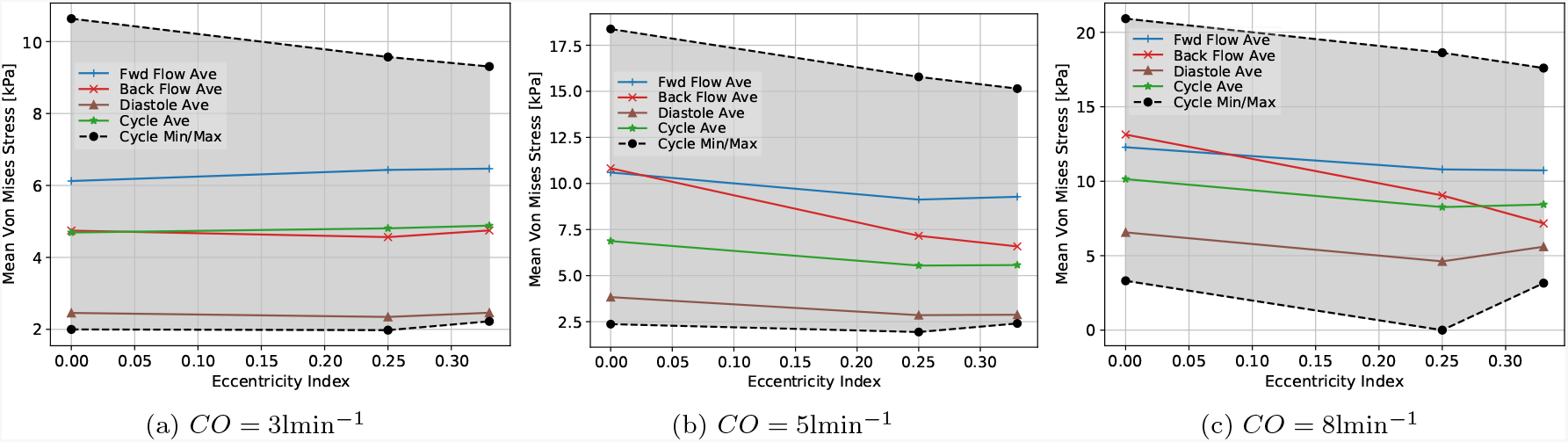
Volume- and phase-averaged von Mises stresses of the valve leaflets as a function of the eccentricity index of the aortic annulus for different cardiac outputs (increasing left to right).

**Figure 26:**
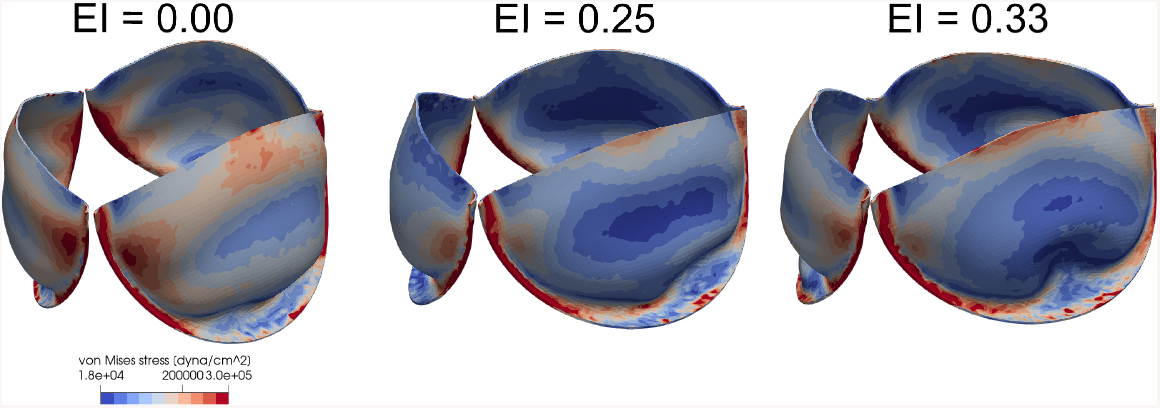
Von Mises stresses in systole for increasing eccentricities (left to right) for conditions corresponding to healthy patient at rest *CO* = 5lmin^−1^.

Figure 25 shows snapshots of the von Mises stresses during systole for the *CO* = 5lmin^−1^ conditions for the different eccentricities. Here it can be seen that while stresses are distributed smoothly for the circular case, they become increasingly concentrated on leaflet edges and attachments for the most eccentric cases. These results show that an increase in eccentricity could be producing a stronger localized concentration of structural stresses, providing hot-points for calcification.

## 4 Conclusions and future work

This *in silico* study demonstrated differences in hydro- and solid-dynamics for a BAVRs of different rigidities and eccentricities. The FSI model was verified and validated against the widely used FSI3 benchmark from Turek & Hron [40]. QoIs defined by the ISO 5840-3 standard, GOA and TPG, showed good agreement with respect to the experimental works of Sigüenza et al. [42] and with respect to the tendencies in Kütting et al. [32]. The effect of both leaflet rigidity and aortic root eccentricity were also evaluated on biomarkers for thrombosis, residence time and shear rate, and for the mechanical leaflet stress, a biomarker for leaflet calcification.

Regarding the leaflet rigidity analysis, it was observed that increasing leaflet Young modulus decreases the EOA while increasing the absolute TPG, exerting an extra workload on the left ventricle. Regions of high thrombogenic risk were identified as those with peak long residence times and high shear rates. It was observed that the predominance of these thrombogenically-risky regions increase for more rigid leaflets. Finally, as leaflet rigidity is increased, peak stresses became more concentrated on both comissures and leaflet attachment points, increasing potential damage at these locations.

The effects of eccentricity on the performance of BAVRs, and their thrombogenic and calcification risks, are analyzed in this work. Tendencies for the orifice area from the reference experiment [32] are reproduced, showing that it is reduced for more eccentric valves in the healthy-patient case for both rest (*CO* = 5lmin^−1^) and exercise (*CO* = 8lmin^−1^) conditions. The tendency of increasing TPG for more eccentric valves is reproduced, indicating a larger load on the left ventricle, potentially increasing risk of LVH and eventually cardiac failure. With respect to thrombogenic risk, as eccentricity is increased, the regions of peak residence time and shear rate are both intensified, increasing the risk of thrombus formation. Finally, it is observed that peak leaflet von Mises stresses increase with stronger eccentricities, enhancing the leaflet calcification risk.

### 4.1 Limitations

The present study carries some limitations. In the cardiac valve validation cases considered here, neither exact geometries nor exact pressure boundary conditions were available, limiting the reproducibility of these experiments [42, 32]. This adds inevitable epistemic errors in this work’s results, particularly in the calculation of the TPGs. A reference experiment with more precisely registered controlled and independent variables would be optimal for a comprehensive validation in order to improve its credibility. Moreover, uncertainty quantification of experimental data would improve the reliability of the numerical-experimental validation. Also regarding boundary conditions in this work, flow rate is imposed at the inlet. This enables computing TPG or GOA as a function of the imposed flow rate curve. Nonetheless, it prohibits computing flow rate as a function of TPG. Imposing pressure at the inlet via a reduced-order compliant model (*ie:* Windkessel model) would enable computing for instance the regurgitant volume, a QoI stated in the ISO 5840-3 standard [41]. Moreover, the material model employed for the leaflet consists of an isotropic Neo-Hookean hyperelastic model. While this material model may be appropriate for the polyurethane valves used by Sigüenza et al. [42] and modeled in section 3.2, the TAVI used in Kütting et al. [32] and modeled in section 3.3 is made out of porcine pericardium, an anisotropic material which may be more accurately modeled by a Fung-type material. Moreover material parameters are not accurately detailed in Sigüenza et al. [42], providing additional potential sources of error. Another limitation of the current work, and most of the available bibliography involving 2-way FSI methods, is that no pre-stresses where considered for the deployed configuration given the oversize with respect to the aortic annulus. Including these would provide more realistic stresses, and furthermore enable quantifying the forces involved between leaflets, TAVI frames and aortic annulus, an important consideration for design and optimization of heart valve prostheses. Nonetheless, as mentioned in Kütting et al. [32], the degree of oversizing used in the experiment was minimal, so that this effect is minimized in the validation of the eccentricity study.

### 4.2 Future work

This work represents the first application of the immersed FSI implementation for deformable solids in *Alya*. The framework presented here is a demonstration of the potential power of this computational tool to evaluate the performance and risks of heart valve devices. The overarching goals of this computational model are two-fold. The first objective is to carry out these evaluations on thousands of bioprostheses designs in populations of virtual patients with a range of comorbidities, while running in short execution times. Second, this same model will be applied in clinical scenarios to aid planning of clinical interventions, by simulating different scenarios before carrying them out on real patients. The next steps towards these objectives are, first of all, to improve the efficiency of the implementation. Second, the accuracy of the current model will be improved by including pre-stresses, anisotropic models and aortic chamber deformation by coupling the immersed FSI with the ALE framework. Third, the model will be introduced in the verification, validation and uncertainty quantification (VVUQ) pipeline developed by the authors’ research group [75] to exhaustively verify and validate this model against *in silico* experiments. Fourth, the heat valve model introduced here will be coupled to the existing fluid-electromechanical model of the human heart implemented in *Alya* [76]. This will enable simulating more complex physiological problems involving tight interaction between the different components of the heart. Finally, the model will be placed in the virtual population interface developed by ELEM Biotech [77], resulting in a service which can be rapidly run on an HPC-cloud for applications in device design, *in silico* clinical trials and clinical planning.

## Acronyms

BSC: Barcelona Supercomputing Center
FSI: Fluid-Structure Interaction
GOA & EOA: Geometric & Effective Orifice Area
TPG: Transvalvular Pressure Gradient
AVR: Aortic Valve Replacement
MAVR, BAVR, TAVR & SAVR: Mechanical, Bioprosthetic, Transcatheter & Surgical Aortic Valve Replacements
CFD: Computational Fluid Dynamics
CSM: Computational Solid Mechanics
QoI: Quantity of Interest
FE: Finite Element
ALE: Arbitrary Lagrangian-Eulerian
BCM & NBCM: Boundary-Conforming & Non Boundary-Conforming Methods
LVH: Left Ventricular Hypertrophy
EI: Eccentricity Index
CO: Cardiac Output
VVUQ: Verification, Validation & Uncertainty Quantification

## Acknowledgments

This project has received funding from the European Union’s Horizon 2020 research and innovation programme under the Marie Skłodowska-Curie grant agreement No. 713673. Authors would like to extend their gratitude to Alfonso Santiago and Vi Vu for the discussions regarding QoIs associated to thrombogenic risk.

## References

[1] Carroll J, Mack M, Vemulapalli S, et al. STS-ACC TVT Registry of Transcatheter Aortic Valve Replacement. J Am Coll Cardiol. 2020; 76(21): 2492–2516. http://dx.doi.org/doi:10.1016/j.jacc.2020.09.595 doi: doi:10.1016/j.jacc.2020.09.595

[2] Vahanian A, Alfieri O, Andreotti F, et al. ESC Committee for Practice Guidelines (CPG); Joint Task Force on the Management of Valvular Heart Disease of the European Society of Cardiology (ESC); European Association for Cardio-Thoracic Surgery (EACTS).. Eur J Cardiothorac Surg. 2012; 42(4): S1–44. http://dx.doi.org/10.1093/ejcts/ezs455 doi: 10.1093/ejcts/ezs455

[3] Nishimura RA, Otto CM, Bonow RO, et al. 2014 AHA/ACC Guideline for the Management of Patients With Valvular Heart Disease: Executive Summary. Circulation 2014; 129(23): 2440–2492. http://dx.doi.org/10.1161/CIR.0000000000000029 doi: 10.1161/CIR.0000000000000029

[4] Chakravarty T, Søndergaard L, Friedman JD, et al. Subclinical leaflet thrombosis in surgical and transcatheter bioprosthetic aortic valves: an observational study. The Lancet 2017; 389: 2383–2392.

[5] Jose J, Sulimov DS, El-Mawardy M, et al. Clinical Bioprosthetic Heart Valve Thrombosis After Transcatheter Aortic Valve Replacement: Incidence, Characteristics, and Treatment Outcomes. JACC: Cardiovascular Interventions 2017; 10(7): 686–697. http://dx.doi.org/https://doi.org/10.1016/j.jcin.2017.01.045 doi: https://doi.org/10.1016/j.jcin.2017.01.045

[6] Mangione FM, Jatene T, Gonçalves A, et al. Leaflet Thrombosis in Surgically Explanted or Post-Mortem TAVR Valves. JACC: Cardiovascular Imaging 2017; 10(1): 82–85. http://dx.doi.org/https://doi.org/10.1016/j.jcmg.2016.11.009 doi: https://doi.org/10.1016/j.jcmg.2016.11.009

[7] Makkar RR, Fontana G, Jilaihawi H, et al. Possible Subclinical Leaflet Thrombosis in Bioprosthetic Aortic Valves. New England Journal of Medicine 2015; 373(21): 2015–2024. PMID: 26436963 http://dx.doi.org/10.1056/NEJMoa1509233 doi: 10.1056/NEJMoa1509233

[8] Makkar RR, Blanke P, Leipsic J, et al. Subclinical Leaflet Thrombosis in Transcatheter and Surgical Bioprosthetic Valves. Journal of the American College of Cardiology 2020; 75(24): 3003–3015. http://dx.doi.org/10.1016/j.jacc.2020.04.043 doi: 10.1016/j.jacc.2020.04.043

[9] Lefkowitz J. Coagulation pathway and physiology: 3–12; ed. K Kottke-Marchang. 2008.

[10] Davie E, Fujikawa K, Kisiel W. The coagulation cascade: initiation, maintenance, and regulation.. Biochemistry 1991; 30(43): 10363–70. http://dx.doi.org/10.1021/bi00107a001 doi: 10.1021/bi00107a001

[11] Cadroy Y, Horbett T, Hanson S. Discrimination between platelet-mediated and coagulation-mediated mechanisms in a model of complex thrombus formation in vivo.. J Lab Clin Med. 1989; 113(4): 436–48.

[12] Casa L, Ku D. Thrombus Formation at High Shear Rates. Annual Review of Biomedical Engineering 2017; 19(1): 415–433. http://dx.doi.org/10.1146/annurev-bioeng-071516-044539 doi: 10.1146/annurev-bioeng-071516-044539

[13] Ducci A, Pirisi F, Tzamtzis S, Burriesci G. Transcatheter aortic valves produce unphysiological flows which may contribute to thromboembolic events: An in-vitro study. Journal of Biomechanics 2016;49: 4080–4089.

[14] Midha PA, Raghav V, Sharma R, et al. The Fluid Mechanics of Transcatheter Heart Valve Leaflet Thrombosis in the Neosinus. Circulation 2017; 136(17): 1598–1609. http://dx.doi.org/10.1161/CIRCULATIONAHA.117.029479 doi: 10.1161/CIRCULATIONAHA.117.029479

[15] Hatoum H, Moore BL, Maureira P, Dollery J, Crestanello J, Dasi LP. Aortic sinus flow stasis likely in valve-in-valve transcatheter aortic valve implantation. The Journal of Thoracic and Cardiovascular Surgery 2017; 154: 32–43.e1.

[16] Vahidkhah K, Barakat M, Abbasi M, et al. Valve thrombosis following transcatheter aortic valve replacement: significance of blood stasis on the leaflets. European Journal of Cardio-Thoracic Surgery 2017; 51(5): 927–935. http://dx.doi.org/10.1093/ejcts/ezw407 doi: 10.1093/ejcts/ezw407

[17] Vahidkhah K, Barakat M, Abbasi M, et al. Valve thrombosis following transcatheter aortic valve replacement: significance of blood stasis on the leaflets. European Journal of Cardio-Thoracic Surgery 2017; 51(5): 927–935. http://dx.doi.org/10.1093/ejcts/ezw407 doi: 10.1093/ejcts/ezw407

[18] Vahidkhah K, Azadani AN. Supra-annular Valve-in-Valve implantation reduces blood stasis on the transcatheter aortic valve leaflets. Journal of Biomechanics 2017; 58: 114–122. http://dx.doi.org/https://doi.org/10.1016/j.jbiomech.2017.04.020 doi: https://doi.org/10.1016/j.jbiomech.2017.04.020

[19] Long CC, Esmaily-Moghadam M, Marsden AL, Bazilevs Y. Computation of Residence Time in the Simulation of Pulsatile Ventricular Assist Devices. Comput. Mech. 2014; 54(4): 911–919. http://dx.doi.org/10.1007/s00466-013-0931-y doi: 10.1007/s00466-013-0931-y

[20] Rossini L, Martinez-Legazpi P, Vu V, et al. A clinical method for mapping and quantifying blood stasis in the left ventricle. Journal of Biomechanics 2016; 49(11): 2152–2161. Selected Articles from the International Conference on CFD in Medicine and Biology (Albufeira, Portugal – August 30th - September 4th, 2015) http://dx.doi.org/https://doi.org/10.1016/j.jbiomech.2015.11.049 doi: https://doi.org/10.1016/j.jbiomech.2015.11.049

[21] Reza MMS, Arzani A. A critical comparison of different residence time measures in aneurysms. Journal of Biomechanics 2019; 88: 122–129. http://dx.doi.org/https://doi.org/10.1016/j.jbiomech.2019.03.028 doi: https://doi.org/10.1016/j.jbiomech.2019.03.028

[22] Plitman Mayo R, Yaakobovich H, Finkelstein A, Shadden SC, Marom G. Numerical models for assessing the risk of leaflet thrombosis post-transcatheter aortic valve-in-valve implantation. Royal Society Open Science 2020; 7(12): 201838. http://dx.doi.org/10.1098/rsos.201838 doi: 10.1098/rsos.201838

[23] Zhu G, Ismail MB, Nakao M, Yuan Q, Yeo JH. Numerical and in-vitro experimental assessment of the performance of a novel designed expanded-polytetrafluoroethylene stentless bi-leaflet valve for aortic valve replacement. PLOS ONE 2019; 14(1): 1–27. http://dx.doi.org/10.1371/journal.pone.0210780 doi: 10.1371/journal.pone.0210780

[24] Deiwick M, Glasmacher B, Baba H, et al. In vitro testing of bioprostheses: influence of mechanical stresses and lipids on calcification.. Ann Thorac Surg. 1983; 66: 206–11. http://dx.doi.org/10.1016/s0003-4975(98)01125-4 doi: 10.1016/s0003-4975(98)01125-4

[25] Thubrikar MJ, Deck JD, Aouad J, Nolan SP. Role of mechanical stress in calcification of aortic bioprosthetic valves.. The Journal of thoracic and cardiovascular surgery 1983; 86 1: 115–25.

[26] Gunning PS, Saikrishnan N, Yoganathan AP, McNamara LM. Total ellipse of the heart valve: the impact of eccentric stent distortion on the regional dynamic deformation of pericardial tissue leaflets of a transcatheter aortic valve replacement. Journal of The Royal Society Interface 2015; 12(113): 20150737. http://dx.doi.org/10.1098/rsif.2015.0737 doi: 10.1098/rsif.2015.0737

[27] Doddamani S, Grushko M, Makaryus Aea. Demonstration of left ventricular outflow tract eccentricity by 64-slice multi-detector CT. Int J Cardiovasc Imaging 2009; 25: 175–181. http://dx.doi.org/https://doi.org/10.1007/s10554-008-9362-9 doi: https://doi.org/10.1007/s10554-008-9362-9

[28] Schultz C, Weustink A, Piazza N, et al. Geometry and Degree of Apposition of the CoreValve ReValving System With Multislice Computed Tomography After Implantation in Patients With Aortic Stenosis. Journal of the American College of Cardiology 2009; 54(10): 911–918. http://dx.doi.org/10.1016/j.jacc.2009.04.075 doi: 10.1016/j.jacc.2009.04.075

[29] Zegdi R, Ciobotaru V, Noghin M, et al. Is It Reasonable to Treat All Calcified Stenotic Aortic Valves With a Valved Stent?: Results From a Human Anatomic Study in Adults. Journal of the American College of Cardiology 2008; 51(5): 579–584. http://dx.doi.org/https://doi.org/10.1016/j.jacc.2007.10.023 doi: https://doi.org/10.1016/j.jacc.2007.10.023

[30] Zegdi R, Lecuyer L, Achouh P, et al. Increased Radial Force Improves Stent Deployment in Tricuspid but Not in Bicuspid Stenotic Native Aortic Valves. The Annals of Thoracic Surgery 2010; 89(3): 768–772. http://dx.doi.org/https://doi.org/10.1016/j.athoracsur.2009.12.022 doi: https://doi.org/10.1016/j.athoracsur.2009.12.022

[31] Wong D, Bertaso A, Liew G, et al. Relationship of aortic annular eccentricity and paravalvular regurgitation post transcatheter aortic valve implantation with CoreValve.. The Journal of invasive cardiology 2013; 25 4: 190–5.

[32] Kuetting M, Sedaghat A, Utzenrath M, et al. In vitro assessment of the influence of aortic annulus ovality on the hydrodynamic performance of self-expanding transcatheter heart valve prostheses. J Biomech. 2014; 47(5): 957–65. http://dx.doi.org/10.1016/j.jbiomech.2014.01.024 doi: 10.1016/j.jbiomech.2014.01.024

[33] Thubrikar M, Piepgrass WC, Shaner TW, Nolan SP. The design of the normal aortic valve. American Journal of Physiology-Heart and Circulatory Physiology 1981; 241(6): H795–H801. PMID: 7325246 http://dx.doi.org/10.1152/ajpheart.1981.241.6.H795 doi: 10.1152/ajpheart.1981.241.6.H795

[34] Salaun E, Zenses AS, Evin M, et al. Effect of oversizing and elliptical shape of aortic annulus on transcatheter valve hemodynamics: An in vitro study. International Journal of Cardiology 2016; 208: 28–35. http://dx.doi.org/https://doi.org/10.1016/j.ijcard.2016.01.048 doi: https://doi.org/10.1016/j.ijcard.2016.01.048

[35] Sun W, Li K, Sirois E. Simulated elliptical bioprosthetic valve deformation: Implications for asymmetric transcatheter valve deployment. Journal of Biomechanics 2010; 43(16): 3085–3090. http://dx.doi.org/https://doi.org/10.1016/j.jbiomech.2010.08.010 doi: https://doi.org/10.1016/j.jbiomech.2010.08.010

[36] Finotello A, Morganti S, Auricchio F. Finite element analysis of TAVI: Impact of native aortic root computational modeling strategies on simulation outcomes. Medical Engineering and Physics 2017; 47: 2–12. http://dx.doi.org/https://doi.org/10.1016/j.medengphy.2017.06.045 doi: https://doi.org/10.1016/j.medengphy.2017.06.045

[37] Sirois E, Mao W, Li K, Calderan J, Sun W. Simulated Transcatheter Aortic Valve Flow: Implications of Elliptical Deployment and Under-Expansion at the Aortic Annulus. Artificial Organs 2018; 42(7): E141–E152. http://dx.doi.org/https://doi.org/10.1111/aor.13107 doi: https://doi.org/10.1111/aor.13107

[38] Viceconti M, Pappalardo F, Rodriguez B, Horner M, Bischoff J, Musuamba Tshinanu F. In silico trials: Verification, validation and uncertainty quantification of predictive models used in the regulatory evaluation of biomedical products. Methods 2021; 185: 120–127. Methods on simulation in biomedicine http://dx.doi.org/https://doi.org/10.1016/j.ymeth.2020.01.011 doi: https://doi.org/10.1016/j.ymeth.2020.01.011

[39] Assessing Credibility of Computational Modeling through Verification and Validation: Application to Medical Devices. standard, ASME; 2018.

[40] Turek S. HJ. Non-Newtonian Rheology in Blood Circulation. arXiv: Fluid Dynamics 2013.

[41] Cardiovascular Implants – Cardiac Valve Prostheses – Part 3: Heart Valve Substitutes Implanted by Transcatheter Techniques - ISO 5840-3:2013(E). standard, International Organization for Standardization; Geneva, CH: 2013.

[42] Sigüenza J, Pott D, Mendez S, et al. Fluid-structure interaction of a pulsatile flow with an aortic valve model: A combined experimental and numerical study. International Journal for Numerical Methods in Biomedical Engineering 2018; 34(4): e2945. e2945cnm.2945 http://dx.doi.org/10.1002/cnm.2945 doi: 10.1002/cnm.2945

[43] Sochi T. Non-Newtonian Rheology in Blood Circulation. arXiv: Fluid Dynamics 2013.

[44] Codina R. Pressure Stability in Fractional Step Finite Element Methods for Incompressible Flows. Journal of Computational Physics 2001; 170(1): 112–140. http://dx.doi.org/https://doi.org/10.1006/jcph.2001.6725 doi: https://doi.org/10.1006/jcph.2001.6725

[45] Lehmkuhl O, Houzeaux G, Owen H, Chrysokentis G, Rodriguez I. A low-dissipation finite element scheme for scale resolving simulations of turbulent flows. Journal of Computational Physics 2019; 390: 51–65. http://dx.doi.org/https://doi.org/10.1016/j.jcp.2019.04.004 doi: https://doi.org/10.1016/j.jcp.2019.04.004

[46] Billiar KL SM. Biaxial mechanical properties of the native and glutaraldehyde-treated aortic valve cusp: Part II–A structural constitutive model.. J Biomech Eng. 2000; 122(4). http://dx.doi.org/10.1115/1.3127261 doi: 10.1115/1.3127261

[47] May-Newman K, Lam C, Yin FCP. A Hyperelastic Constitutive Law for Aortic Valve Tissue. Journal of Biomechanical Engineering 2009; 131(8). 081009 http://dx.doi.org/10.1115/1.3127261 doi: 10.1115/1.3127261

[48] Donea J, Huerta A, Ponthot JP, Rodríguez-Ferran A. Arbitrary Lagrangian–Eulerian Methodsch. 14; John Wiley & Sons, Ltd. 2004

[49] Peskin CS. Flow patterns around heart valves: A numerical method. Journal of Computational Physics 1972; 10(2): 252–271. http://dx.doi.org/https://doi.org/10.1016/0021-9991(72)90065-4 doi: https://doi.org/10.1016/0021-9991(72)90065-4

[50] Wang X, Zhang L. Interpolation functions in the immersed boundary and finite element methods. Computational Mechanics 2010; 45: 321–334. http://dx.doi.org/10.1007/s00466-009-0449-5 doi: 10.1007/s00466-009-0449-5

[51] Vázquez M, Houzeaux G, Koric S, et al. Alya: Multiphysics engineering simulation toward exascale. Journal of Computational Science 2016; 14: 15–27. The Route to Exascale: Novel Mathematical Methods, Scalable Algorithms and Computational Science Skills http://dx.doi.org/https://doi.org/10.1016/j.jocs.2015.12.007 doi: https://doi.org/10.1016/j.jocs.2015.12.007

[52] Mira D, Zavala-Ake M, Avila Mea. Heat Transfer Effects on a Fully Premixed Methane Impinging Flame. Flow Turbulence Combust 2016; 97: 339–36. http://dx.doi.org/https://doi.org/10.1007/s10494-015-9694-1 doi: https://doi.org/10.1007/s10494-015-9694-1

[53] Calmet H, Gambaruto AM, Bates AJ, Vázquez M, Houzeaux G, Doorly DJ. Large-scale CFD simulations of the transitional and turbulent regime for the large human airways during rapid inhalation. Computers in Biology and Medicine 2016; 69: 166–180. http://dx.doi.org/https://doi.org/10.1016/j.compbiomed.2015.12.003 doi: https://doi.org/10.1016/j.compbiomed.2015.12.003

[54] Gövert S, Mira D, Zavala-Ake M, Kok J, Vázquez M, Houzeaux G. Heat loss prediction of a confined premixed jet flame using a conjugate heat transfer approach. International Journal of Heat and Mass Transfer 2017; 107: 882–894. http://dx.doi.org/https://doi.org/10.1016/j.ijheatmasstransfer.2016.10.122 doi: https://doi.org/10.1016/j.ijheatmasstransfer.2016.10.122

[55] Rodriguez I, Lehmkuhl O, Soria M, Gómez S, Domínguez-Pumar M, Kowalski L. Fluid dynamics and heat transfer in the wake of a sphere. International Journal of Heat and Fluid Flow 2019; 76: 141–153. http://dx.doi.org/https://doi.org/10.1016/j.ijheatfluidflow.2019.02.004 doi: https://doi.org/10.1016/j.ijheatfluidflow.2019.02.004

[56] Santiago A, Zavala-Aké M, Borell R, Houzeaux G, Vázquez M. HPC compact quasi-Newton algorithm for interface problems. Journal of Fluids and Structures 2020; 96: 103009. http://dx.doi.org/10.1016/j.jfluidstructs.2020.103009 doi: 10.1016/j.jfluidstructs.2020.103009

[57] Oyarzun G, Mira D, Houzeaux G. Performance assessment of CUDA and OpenACC in large scale combustion simulations. 2021.

[58] Dunne T, Rannacher R. Adaptive Finite Element Approximation of Fluid-Structure Interaction Based on an Eulerian Variational Formulation. In: Bungartz HJ, Schäfer M., eds. Fluid-Structure Interaction Springer Berlin Heidelberg; 2006; Berlin, Heidelberg: 110–145.

[59] Heil M, Hazel AL, Boyle J. Solvers for large-displacement fluid–structure interaction problems: segregated versus monolithic approaches. Computational Mechanics 2008; 43: 91–101.

[60] Sun X, Steve Suh C, Sun C, Yu B. Vortex-induced vibration of a flexible splitter plate attached to a square cylinder in laminar flow. Journal of Fluids and Structures 2021; 101: 103206. http://dx.doi.org/https://doi.org/10.1016/j.jfluidstructs.2020.103206 doi: https://doi.org/10.1016/j.jfluidstructs.2020.103206

[61] Breuer M, De Nayer G, Münsch M, Gallinger T, Wüchner R. Fluid-structure interaction using a partitioned semi-implicit predictor-corrector coupling scheme for the application of large-eddy simulation. Journal of Fluids and Structures 2012; 29: 107–130. http://dx.doi.org/10.1016/j.jfluidstructs.2011.09.003 doi: 10.1016/j.jfluidstructs.2011.09.003

[62] Bhardwaj R, Mittal R. Benchmarking a Coupled Immersed-Boundary-Finite-Element Solver for Large-Scale Flow-Induced Deformation. AIAA Journal 2012; 50: 1638–1642.

[63] Tian FB, Dai H, Luo H, Doyle JF, Rousseau B. Fluid–structure interaction involving large deformations: 3D simulations and applications to biological systems. Journal of Computational Physics 2014; 258: 451–469. http://dx.doi.org/https://doi.org/10.1016/j.jcp.2013.10.047 doi: https://doi.org/10.1016/j.jcp.2013.10.047

[64] E. Griffith B, Luo X. Hybrid finite difference/finite element immersed boundary method. International Journal for Numerical Methods in Biomedical Engineering 2017; 33(12): e2888. e2888 cnm.2888 http://dx.doi.org/10.1002/cnm.2888 doi: 10.1002/cnm.2888

[65] Westerhof N, Lankhaar J, Westerhof B. The arterial Windkessel.. Med Biol Eng Comput 2009; 47: 131–131. http://dx.doi.org/https://doi.org/10.1007/s11517-008-0359-2 doi: https://doi.org/10.1007/s11517-008-0359-2

[66] Lindman B, Clavel MA, Mathieu P, et al. Calcific aortic stenosis. Nature Reviews Disease Primers 2016; 2: 16006. http://dx.doi.org/10.1038/nrdp.2016.6 doi: 10.1038/nrdp.2016.6

[67] Rayz V, Boussel L, Ge L, et al. Flow residence time and regions of intraluminal thrombus deposition in intracranial aneurysms.. Ann Biomed Eng. 2010; 38(10): 3058–69. http://dx.doi.org/10.1007/s10439-010-0065-8 doi: 10.1007/s10439-010-0065-8

[68] Gorbet MB, Sefton MV. Biomaterial-associated thrombosis: roles of coagulation factors, complement, platelets and leukocytes. Biomaterials 2004; 25(26): 5681–5703. http://dx.doi.org/https://doi.org/10.1016/j.biomaterials.2004.01.023 doi: https://doi.org/10.1016/j.biomaterials.2004.01.023

[69] Rukhlenko OS, Dudchenko OA, Zlobina KE, Guria GT. Mathematical Modeling of Intravascular Blood Coagulation under Wall Shear Stress. PLOS ONE 2015; 10(7): 1–16. http://dx.doi.org/10.1371/journal.pone.0134028 doi: 10.1371/journal.pone.0134028

[70] Schofield Z, Baksamawi H, Campos J, et al. The role of venous stiffness in the insurgence of deep vein thrombosis. Communications Materials 2020; 1: 1234567890. http://dx.doi.org/10.1038/s43246-020-00066-2 doi: 10.1038/s43246-020-00066-2

[71] Dweck MR, Boon NA, Newby DE. Calcific aortic stenosis: a disease of the valve and the myocardium. Journal of the American College of Cardiology 2012; 60(19): 1854–1863.

[72] Hein S, Arnon E, Kostin S, et al. Progression from compensated hypertrophy to failure in the pressure-overloaded human heart: structural deterioration and compensatory mechanisms. Circulation 2003; 107(7): 984–991.

[73] Chin CW, Vassiliou V, Jenkins WS, Prasad SK, Newby DE, Dweck MR. Markers of left ventricular decompensation in aortic stenosis. Expert review of cardiovascular therapy 2014; 12(7): 901–912.

[74] De Marchena E, Mesa J, Pomenti S, et al. Thrombus Formation Following Transcatheter Aortic Valve Replacement. JACC: Cardiovascular Interventions 2015; 8(5): 728–739. TAVR Focus Issue - http://dx.doi.org/https://doi.org/10.1016/j.jcin.2015.03.005 doi: https://doi.org/10.1016/j.jcin.2015.03.005

[75] Santiago A, Butakoff C, Eguzkitza B, et al. Design and execution of a Verification, Validation, and Uncertainty Quantification plan for a numerical model of left ventricular flow after LVAD implantation. 2021

[76] Santiago A, Aguado-Sierra J, Miguel ZA, al. e. Fully coupled fluid-electro-mechanical model of the human heart for supercomputers. Int J Numer Meth Biomed Engng. 2018; 34: 3140. http://dx.doi.org/https://doi.org/10.1002/cnm.3140 doi: https://doi.org/10.1002/cnm.3140

[77] Aguado-Sierra J, Butakoff C, Brigham R, et al. In-silico clinical trial using high performance computational modeling of a virtual human cardiac population to assess drug-induced arrhythmic risk. medRxiv 2021. http://dx.doi.org/10.1101/2021.04.21.21255870 doi: 10.1101/2021.04.21.21255870

